# Exploring functional connectivity in clinical and data-driven groups of preterm and term adults

**DOI:** 10.1101/2024.01.22.576651

**Authors:** Laila Hadaya, František Váša, Konstantina Dimitrakopoulou, Mansoor Saqi, Sukhwinder S Shergill, David A Edwards, Dafnis Batalle, Robert Leech, Chiara Nosarti

## Abstract

**Background:** Adults born very preterm (i.e., at <33 weeks’ gestation) are more susceptible to long-lasting structural and functional brain alterations and cognitive and socio-emotional difficulties, compared to full-term controls. However, behavioural heterogeneity within very preterm and full-term individuals makes it challenging to find biomarkers of specific outcomes. To address these questions, we parsed brain-behaviour heterogeneity in participants subdivided according to their clinical birth status (very preterm vs full-term) and/or data-driven behavioural phenotype (regardless of birth status).

**Methods:** The Network Based Statistic approach was used to identify topological components of resting state functional connectivity differentiating between i) 116 very preterm and 83 full-term adults (43% and 57% female, respectively), and ii) data-driven behavioural subgroups identified using consensus clustering (n= 156, 46% female). Age, sex, socio-economic status, and in-scanner head motion were used as confounders in all analyses. Post-hoc two-way group interactions between clinical birth status and behavioural data-driven subgrouping classification labels explored whether functional connectivity differences between very preterm and full-term adults varied according to distinct behavioural outcomes.

**Results:** Very preterm compared to full-term adults had poorer scores in selective measures of cognitive and socio-emotional processing and displayed complex patterns of hyper- and hypo-connectivity in subsections of the default mode, visual, and ventral attention networks. Stratifying the study participants in terms of their behavioural profiles (irrespective of birth status), identified two data-driven subgroups: An “*At-risk”* subgroup, characterised by increased cognitive, mental health, and socio-emotional difficulties, displaying hypo-connectivity anchored in frontal opercular and insular regions, relative to a *“Resilient”* subgroup with more favourable outcomes. No significant interaction was noted between clinical birth status and behavioural data-driven subgrouping classification labels in terms of functional connectivity.

**Conclusions:** Functional connectivity differentiating between very preterm and full-term adults was dissimilar to functional connectivity differentiating between the data-driven behavioural subgroups. We speculate that functional connectivity alterations observed in very preterm relative to full-term adults may confer both risk and resilience to developing behavioural sequelae associated with very preterm birth, while the localised functional connectivity alterations seen in the “*At-risk”* subgroup relative to the “*Resilient*” subgroup may underlie less favourable behavioural outcomes in adulthood, irrespective of birth status.

## 1.1 Introduction

Very preterm birth (VPT; i.e., at <33 weeks’ gestation) occurs during a rapid stage of brain development, making those born VPT vulnerable to neurological insult (Volpe, 2009) and long-lasting difficulties in attention, executive function, and socio-emotional processing (Anderson et al., 2021; Johnson and Marlow, 2011; Kroll et al., 2017). Functional connectivity alterations in brain regions and networks important for cognitive and affective processing have also been reported in VPT samples across the lifespan, and have been studied amongst the possible biological mechanisms underlying the behavioural difficulties associated with VPT birth (Bäuml et al., 2015; Kanel et al., 2022; Mueller et al., 2022; Papini et al., 2016; Ramphal et al., 2020; Rogers et al., 2017; Siffredi et al., 2022; Sylvester et al., 2018). It is important to highlight, however, that not only have previous studies identified brain changes associated with behavioural difficulties in those born VPT, but have also characterised neural adaptions which support domain-specific performance (Daamen et al., 2014; Finke et al., 2015; Nosarti et al., 2006, 2009; Schafer et al., 2009). These findings, therefore, indicate that the functional reorganisation of the VPT brain has complex implications for outcomes, as it may probe both risk and resilience to behavioural difficulties.

Further complicating the understanding of brain-behavioural relationships in VPT populations, is the fact that those born preterm tend to exhibit heterogenous behavioural outcomes. Previous studies aiming to stratify this heterogeneity implemented latent profile analyses using behavioural measures from both preterm and FT born children (Burnett et al., 2019; Johnson et al., 2018; Lean et al., 2020). Their results indicated that while those born preterm were more likely to present with psychiatric, cognitive, or socio-emotional difficulties, some preterm children displayed distinct profiles characterised by fewer or no behavioural difficulties. Moreover, while FT children predominantly exhibited more normo-typical behavioural profiles, some FT children displayed behavioural difficulties similar to those observed in preterm children (Burnett et al., 2019; Johnson et al., 2018; Lean et al., 2020). Together, these findings indicate that VPT and FT groups exhibit both within- and between-group heterogeneity, which needs to be addressed in order to develop individually tailored and biologically specific interventions aimed at supporting healthy development (Cuthbert and Insel, 2013; Morris et al., 2022). This can be achieved by, firstly, implementing data-driven stratification approaches to identify distinct subgroups of individuals exhibiting similar behavioural profiles, irrespective of their birth status, and secondly, by investigating brain correlates differentiating between the distinct data-driven behavioural subgroups.

Similarly, individuals belonging to distinct diagnostic and non-diagnostic psychiatric groups also exhibit within- and between-group heterogeneity in terms of phenotypic profiles. Recent studies implementing such approaches in psychiatric populations have successfully identified patterns of structural and functional connectivity characterising distinct data-driven behavioural subgroups irrespective of diagnostic labels (Astle et al., 2019; Bathelt et al., 2018; Jones et al., 2021; Mareva et al., 2023; Siugzdaite et al., 2020; Vandewouw et al., 2023). A small number of studies in VPT children followed similar methodological approaches and investigated the underlying brain changes differentiating within-group behavioural heterogeneity. Results of these studies showed that early brain insult (Bogičević et al., 2021; Ross et al., 2016) and structural and functional brain alterations (Hadaya et al., 2023; Lean et al., 2020) characterised the distinct subgroups. However, it remains to be explored whether the heterogeneity in behavioural outcomes seen within and between VPT and FT born individuals persists into adulthood, and if it does, whether resting state functional connectivity (rsFC) changes may be associated with distinct data-driven behavioural phenotypes, irrespective of gestational age at birth.

Our study firstly aimed to identify long-lasting neurodevelopmental alterations associated with VPT birth, by investigating differences in rsFC and behavioural outcomes between VPT and FT born adults. Secondly, our study aimed to delineate behavioural heterogeneity in VPT and FT born adults irrespective of gestational age at birth, by using a robust data-driven consensus clustering approach to stratify participants based on behavioural measures (executive function, attention, intelligence, socio-emotional processing, psychopathology, and autistic traits), and to explore whether resultant data-driven behavioural subgroups would exhibit differences in rsFC. Finally, post-hoc two-way group interactions between clinical (i.e., VPT vs FT birth) and behavioural (i.e., data-driven subgrouping) classification labels were used to explore whether rsFC pattern differences between VPT and FT adults, varied according to distinct behavioural outcomes.

## 1.2 Methods

### 1.2.1 Study design

#### Participants

VPT infants (i.e., born at <33 weeks of gestation) were recruited at birth from the Neonatal Unit at University College London Hospital (London, UK) between 1979 and 1985. Enrolled participants received cranial ultrasonographic imaging several times during the first week of life and weekly until discharge from hospital (Stewart et al., 1983) and were subsequently followed up in childhood at 1, 4 and 8 years of age (Roth et al., 1994; Stewart et al., 1989), adolescence (15 years), early (20 years), and middle adulthood (30 years) (Karolis et al., 2017). Age-matched controls, born at FT (37-42 weeks of gestation), were recruited from the community in middle adulthood. Exclusion criteria for the controls were any clinical complications at birth (i.e., prolonged gestation at >42 weeks, low birth weight <2500g, receiving endotracheal mechanical ventilation). Exclusion criteria for both VPT and FT participants included severe hearing and motor impairments, or history of neurological complications (i.e., meningitis, head injury, cerebral infections). For this study, we used neuroimaging and behavioural data from the middle adulthood follow-up.

Research study practices were conducted in accordance with the Declaration of Helsinki. Ethical approval was granted by the South London and Maudsley Research and Ethics Committee and the Psychiatry, Nursing and Midwifery Research Ethics Subcommittee (PNM/12/13-10), King’s College London. All participants were native English speakers. Written informed consent was obtained from all study participants and participant privacy rights were observed.

#### Clinical and socio-demographic details

Gestational age at birth and birth weight were collected from medical discharge notes for VPT participants. Participants born VPT were classified into three groups, according to cranial ultrasound diagnosis: no evidence of perinatal brain injury (no injury), grade I – II periventricular haemorrhage without ventricular dilation (minor injury) and grade III – IV periventricular haemorrhage with ventricular dilation (major injury) (Nosarti et al., 2002).

For both VPT and FT groups, self-reported ethnicity was recorded according to the following groups: African, Afro-Caribbean, Caucasian/White, Indian Subcontinent, and Other. Socio-economic status was defined according to participants’ self-reported occupation at the time of the study and parental occupation at birth. Occupations were categorised according to the Office of National Statistics, 1980 Standard Occupation Classification; I: Higher managerial, administrative, and professional occupations; II: Intermediate occupations, small employers, and own account workers; III: Routine and manual occupations – lower supervisory and technical and semi-routine and routine occupations.

#### Cognitive assessments

The following cognitive assessments were administered to measure language, executive attention, and general intelligence: Hayling Sentence Completion Test (HSCT) (Burgess and Shallice, 1997); Controlled Oral Word Association Test (COWAT-FAS) (Benton et al., 1983); four subtests from the Cambridge Neurophysiological Test Automated Battery (CANTAB) 2003 eclipse version (Fray et al., 1996): 1) Stockings of Cambridge (SOC), 2) Intra-Extra Dimensional Set Shift (IED), 3) Paired Associates Learning (PAL), and 4) Motor Screening Task (MOT); the Trail Making Task – B (TMT-B) (Tombaugh, 2004); Continuous Performance Test – 2^nd^ edition (CPT) (Conners et al., 2003); and Wechsler Abbreviated Scale of Intelligence (WASI) (Wechsler, 1999). Specific task descriptions are detailed in Table SM 1.

#### Psychiatric and behavioural assessments

General psychopathology was measured using the Comprehensive Assessment of At-Risk Mental States (CAARMS) (Yung et al., 2005), a semi-structured clinical interview which measures aspects of psychopathology relating to mania, depression, suicidality and self-harm, mood swings/lability, anxiety, obsessive compulsive disorder symptoms, dissociative symptoms, and impaired tolerance to normal stress; scores on the general psychopathology subscale were used in our analyses. The self-administered General Health Questionnaire (GHQ-12) (Goldberg and Williams, 1991) was used to measure general well-being, Peters Delusion Inventory (PDI) (Peters et al., 2004) to measure delusional ideation traits, Autism Quotient (AQ-10) (Allison et al., 2012; Booth et al., 2013) to measure autism traits (i.e., social interaction, communication, attention switching, attention to detail, and imagination), Social Adjustment Scale (SAS) (Weissman and Bothwell, 1976) to measure participants’ satisfaction with their social situation, and Role Functioning Scale (RFS) (Goodman et al., 1993) to measure individuals’ ability to function in their daily life. The Emotion recognition task (ERT) (Montagne et al., 2007) was administered to measure participants’ ability to recognise expressed emotions (happiness, sadness, surprise, anger, disgust and fear), as described in our previous work (Papini et al., 2016).

#### Structural and functional magnetic resonance imaging (MRI) acquisition

MRI data were acquired at the Maudsley Hospital, London, using a 3 Tesla Signa MR scanner (General Electric Healthcare). Structural fast spoiled gradient-echo (FSPGR) pulse sequence T1-weighted images were collected using the following sequence parameters: TR=7.1 ms, TE=2.8 ms, matrix=256×256, voxel size=1.1 mm isotropic. Gradient echo EPI resting state functional MRI data were collected while participants stared at a central cross on a screen for 8 minutes 32 s, using the following parameters 256 volumes, TR=2000 ms, TE=30 ms, flip angle=75 degrees, matrix=64×64, 37 non-contiguous slices of 2.4 mm thickness, 1.1 mm interslice gap, and 3.4 mm in-plane resolution.

### 1.2.2 MRI data pre-processing

Resting state functional MRI data pre-processing was performed using fMRIPrep 20.1.1, RRID:SCR_016216 (Esteban et al., 2019), which is based on Nipype 1.5.0, RRID:SCR_002502 (Gorgolewski et al., 2011). In summary, steps included skull stripping, slice-time correction, co-registration to the T1w reference image using boundary-based registration (Greve and Fischl, 2009) and head motion estimation (i.e., global signal and six motion parameters: three translation and three rotation parameters). The complete pre-processing protocol is detailed in the Supplementary Material.

After pre-processing, data were de-noised by regressing out estimated motion confounders (i.e., global signal and six motion parameters: three translation and three rotation parameters) using the FMRIB Software Library (FSL) *fsl_regfilt* command (Jenkinson et al., 2012). A band-pass filter (0.01 – 0.1 Hz) was applied to the data using the AFNI software *3dBandpass* command (Cox, 1996). Participants were excluded if they exhibited excessive in-scanner head motion (i.e., mean frame-wise displacement (FD) exceeding 0.4mm or a maximum FD exceeding 4mm) or had functional MRI scans showing poor alignment with anatomical data. Sample sizes and participant exclusions are summarised in a flowchart in Figure SM 1.

### 1.2.3 Brain parcellation and rsFC estimation

Resting state functional MRI data were parcellated into bilaterally symmetric cortical regions using the Human Connectome Project Multi-Modal Parcellation; HCP-MMP (v1) atlas (Glasser et al., 2016) and bilateral subcortical FreeSurfer regions (Fischl, 2012). The two bilateral hippocampal regions from the HCP-MMP atlas were excluded as these regions were included as part of the FreeSurfer subcortical segmentation, resulting in a total of 374 regions included in our analyses (i.e., 358 HCP-MMP atlas bilateral cortical regions and 16 FreeSurfer bilateral subcortical regions).

An average of the functional MRI blood oxygen level-dependent (BOLD) signal time series across all voxels in each parcellation was used to estimate the regional time series for each of the 374 brain regions. For each participant, rsFC matrices were calculated using Pearson’s correlations between pairs of all 374 regional time series. A threshold of 0.2 was used to eliminate weak correlations (i.e., weights of edges with r^2^ 0.2 were retained) and a Fisher Z-transformation was applied (Buckner et al., 2009; Fenn-Moltu et al., 2022; Zalesky et al., 2016).

### 1.2.4 Consensus clustering

To partition participants (both VPT and FT; n=156) into data-driven behavioural subgroups, a consensus clustering pipeline (Figure 1) was implemented using the following 13 behavioural measures as input features: COWAT-FAS mean total words produced, SOC total number of problems solved, IED total errors adjusted, MOT mean reaction time, TMT-B time elapsed, CPT total reaction time, full-scale IQ, total PDI score, total AQ10 score, CAARMS total general psychopathology score, total GHQ score, ERT total number of correct responses and total SAS score (see Supplementary Material for data pre-processing and feature selection procedures).

**Figure 1.**
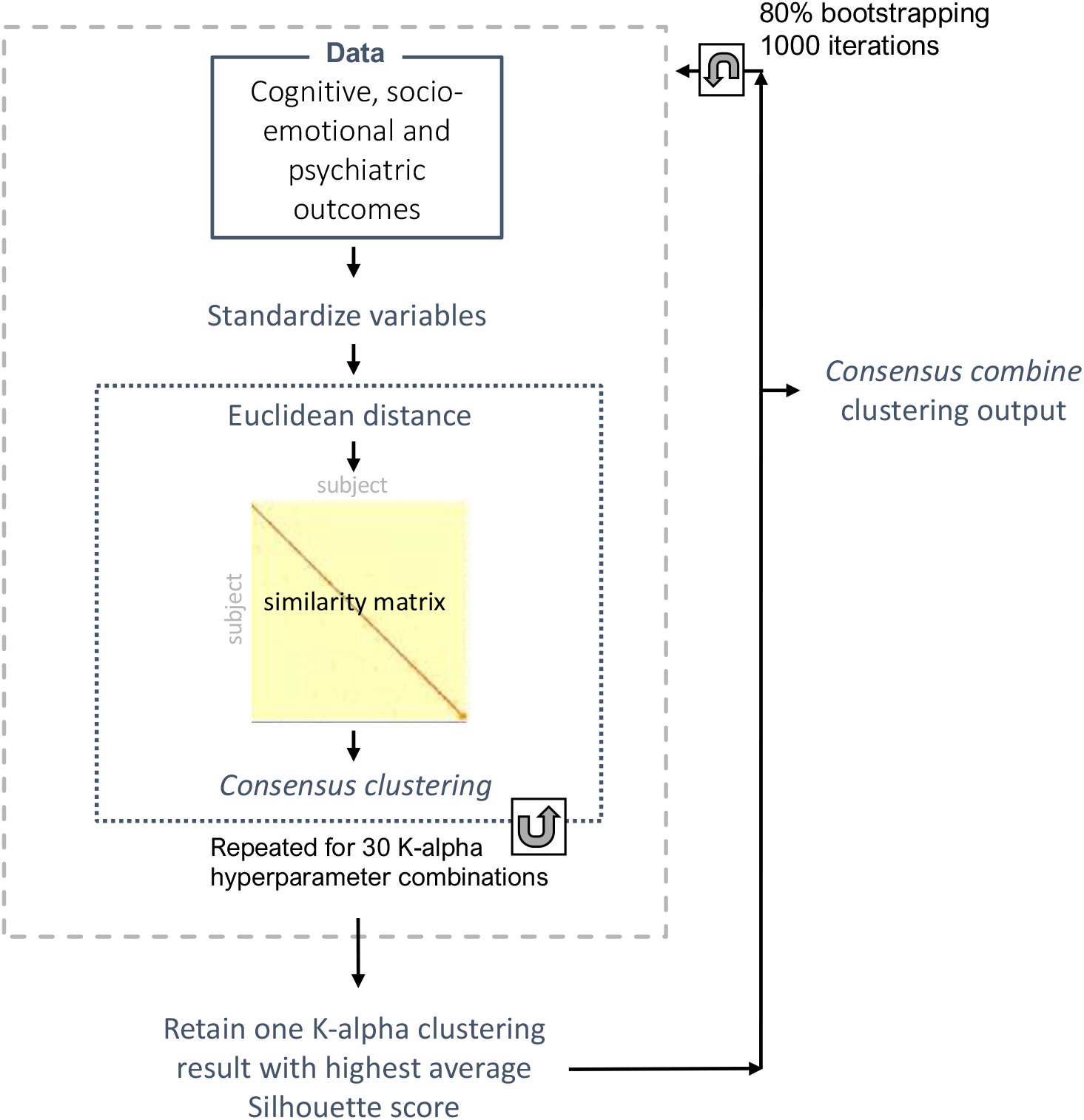
Consensus clustering pipeline followed.

Each variable was first standardised to have a mean = 0 and standard deviation = 1, and an Euclidean distance matrix of the input data was calculated. A similarity matrix (network) was then calculated from the distance matrix, using the *affinityMatrix* function (SNFtool R package) (Wang et al., 2018), which utilises two hyperparameters: neighbourhood size (K) and alpha (edge weighting parameter) that help increase the signal to noise ratio and in turn improve result validity and reliability. K corresponds to the number of surrounding nodes to consider for each node in the similarity network and alpha determines a threshold for the strength of the edges in the similarity network (i.e., pairwise similarity between nodes within the sample). Greater K values result in more dense similarity networks and smaller values result in more sparse similarity networks, while greater alpha values result in weaker edges being retained and smaller alpha values result in similarity networks which retain edges with higher similarity. Thirty different K-alpha combinations were used to generate thirty similarity networks based on the following values: K = 10, 15, 20, 25, 30 and alpha = 0.3, 0.4, 0.5, 0.6, 0.7, 0.8. These values lie within the ranges recommended in the SNFtool package: 10–30 for K and 0.3–0.8 alpha (Wang et al., 2018). Each of the resultant thirty similarity networks was successively inputted into the consensus clustering algorithm (*ConsensusClusterPlus* function, ConsensusClusterPlus R package) (Wilkerson and Hayes, 2010) which performs agglomerative hierarchical clustering following a nested bootstrapping (n=1000) spectral clustering for each of the thirty similarity networks. From the thirty resultant clustering outputs, the solution with the highest average silhouette width score was retained.

In order to improve the generalisability of our solution and avoid overfitting of hyperparameter selection, the steps described in the above paragraph were repeated 1,000 times where a randomised selection of 80% of the sample was used each time. The final resultant 1,000 clustering outputs were then fed into a hierarchical clustering function (*consensus_combine,* DiceR package) (Chiu and Talhouk, 2018), to output a final consensus clustering result based on the consensus matrix.

To determine the optimal number of clusters (C), Eigengap and Rotation Cost metrics were firstly used to estimate the best and second-best number of clusters (*estimateNumberOfClustersGivenGraph* function SNFtool R package) (Wang et al., 2018) for each of the thirty K-alpha combinations, identifying C=2, C=3, and C=5 as the top three clustering solutions. We then ran the described consensus clustering pipeline three separate times, once for each of these solutions (C=2, C=3, and C=5), and subsequently calculated consensus matrices and silhouette scores for each cluster solution. Resultant consensus matrix and silhouette score outputs suggested an optimal number of clusters of C=2 (Figure SM 2); therefore, we evaluate subgroups obtained from the C=2 solution.

The consensus clustering pipeline implemented here is adapted from the integrative clustering method used in our previous work (Hadaya et al., 2023), code: https://github.com/lailahadaya/preterm-ExecuteSNF.C), where we do not apply the data-integration step in the current study.

### 1.2.5 Statistical analyses

#### 1.2.5.1 Evaluation of clinical, socio-demographic, and behavioural profiles

The non-parametric Wilcox Rank Sum T-test was used for continuous variables and Chi-squared or Fischer’s Exact tests for categorical variables. Effect sizes were calculated using Wilcoxon Glass Rank Biserial Correlation for continuous variables and Cramer’s V (V) for categorical variables. False Discovery Rate (FDR) was used to account for multiple comparison testing (Benjamini and Hochberg, 1995). Sensitivity analyses using non-parametric permutation testing (5000 permutations) adjusted for potential covariates (age, sex, and socio-economic status) (França et al., 2022). P-values<0.05 were considered to be statistically significant.

#### 1.2.5.2 Between-group differences in rsFC at a topological network-level

The Network Based Statistic (NBS), a cluster-based statistics approach, was applied (Zalesky et al., 2010). NBS implements the following steps: 1) mass-univariate testing with a suitable statistical test of interest on all possible connections (i.e., edges), 2) next, only edges with p-values below a pre-defined threshold (p-NBS-Threshold) are maintained, 3) retained suprathreshold edges are then used to identify topologically connected structures (referred to as NBS ‘components’) present amongst the collection of suprathreshold edges using breadth-first search (Ahuja et al., 1993), and finally, 4) permutation testing is used to assign a Family Wise Error Rate corrected p-value (p-FWER) for each identified component, based on the component’s strength. NBS testing is derived from traditional cluster-based thresholding of statistical maps; however, rather than generating clusters of voxels with spatial proximity in physical space, NBS can be applied to graph-like structures to generate clusters with interconnected edges in topological space (Nichols and Holmes, 2002; Zalesky et al., 2010). An advantage of using NBS, compared to an approach that controls for FWER at an edgewise basis (such as False-Discovery Rate), is that it can provide increased statistical power by detecting the effect of interest in a collection of connections which are collectively contributing to the effect of interest as opposed to uniquely contributing to the effect on an individual edgewise-level.

Selecting a threshold in NBS (described in step 2 above) is a relatively arbitrary choice, which can be determined by experimenting with a selection of conservative and stringent thresholds (Zalesky et al., 2010). We ran NBS testing at three different p-value thresholds (i.e., p-NBS-Threshold = 0.05, 0.01, and 0.001) to identify relevant suprathreshold edges to be grouped into NBS components for further analysis. We implemented NBS testing with 1000 permutations using the NBR R package *nbr_lm* function (NBR) (Gracia-Tabuenca and Alcauter, 2020). Statistical models tested included the following covariates: mean FD (as a measure of in-scanner head motion), sex, age, and socio-economic status. The same sets of methods were implemented to identify differences in rsFC between 1) VPT and FT individuals and 2) data-driven behavioural subgroups.

NBS generates two resultant outputs: 1) component strength or intensity – i.e., the sum of test statistic (T-statistic) values from all edges within the significant component, and 2) component size or extent – i.e., the number of connections comprising the significant component. We also calculated the number of connections belonging to each node within the component as a proportion of the total number of possible edges within that component and presented results graphically using the ggseg3d R package (Mowinckel and Vidal-Piñeiro, 2020). To measure within and between network connectivity, we labelled nodes according to seven previously defined intrinsic connectivity networks (i.e., visual, somatomotor, dorsal attention, ventral attention (VAN), limbic, frontoparietal, and default mode (DMN) networks) (Yeo et al., 2011) and an eighth network comprised of 16 subcortical regions (Váša et al., 2020) and calculated connectivity proportion and strength; code accessible at: https://github.com/frantisekvasa/functional_network_development/blob/master/nspn.fmri.R.

### 1.2.6 Post-hoc exploratory analyses

We estimated the extent of nodal and edgewise overlap between the NBS components characterising clinical (i.e., VPT vs FT birth) and data-driven behavioural subgrouping classifications using the Sørensen-Dice similarity coefficient, which is calculated as the ratio of two times the number of overlapping features between two sets, over the total number of features present across both sets (Sørensen, 1948), with values ranging between 0 and 1.

Post-hoc exploratory NBS analyses investigated whether differences in rsFC between VPT and FT clinical groups varied according to distinct behavioural outcomes, using two-way group interactions between clinical and data-driven behavioural classification labels.

We also investigated differences in early clinical risk (i.e., gestational age at birth, birth weight, and perinatal brain injury) and socio-demographic measures between VPT adults belonging to the distinct data-driven behavioural subgroups, and in socio-demographic measures between FT adults in the distinct data-driven subgroups.

## 1.3 Results

### 1.3.1 VPT and FT groups

The socio-demographic and clinical profiles of VPT and FT adults are summarised in Table 1 and their behavioural outcomes in Table 2 and Figure 2A. In summary, adults born VPT had lower full-scale IQ (WASI), attention set shifting (CANTAB-IED), and emotion recognition (ERT) scores than adults born FT. Head motion during functional MRI acquisition was greater in the VPT (median FD = 0.15mm, range=0.07 – 0.40) than the FT group (median FD=0.12mm, range=0.05 – 0.35; p<0.001). Supplementary analyses show that VPT adults excluded from analyses (n=37) for reasons described in Figure SM 1, had relatively poorer cognitive and socio-emotional scores relative to those VPT adults included in the analyses (n=116) (Table SM 2).

**Figure 2.**
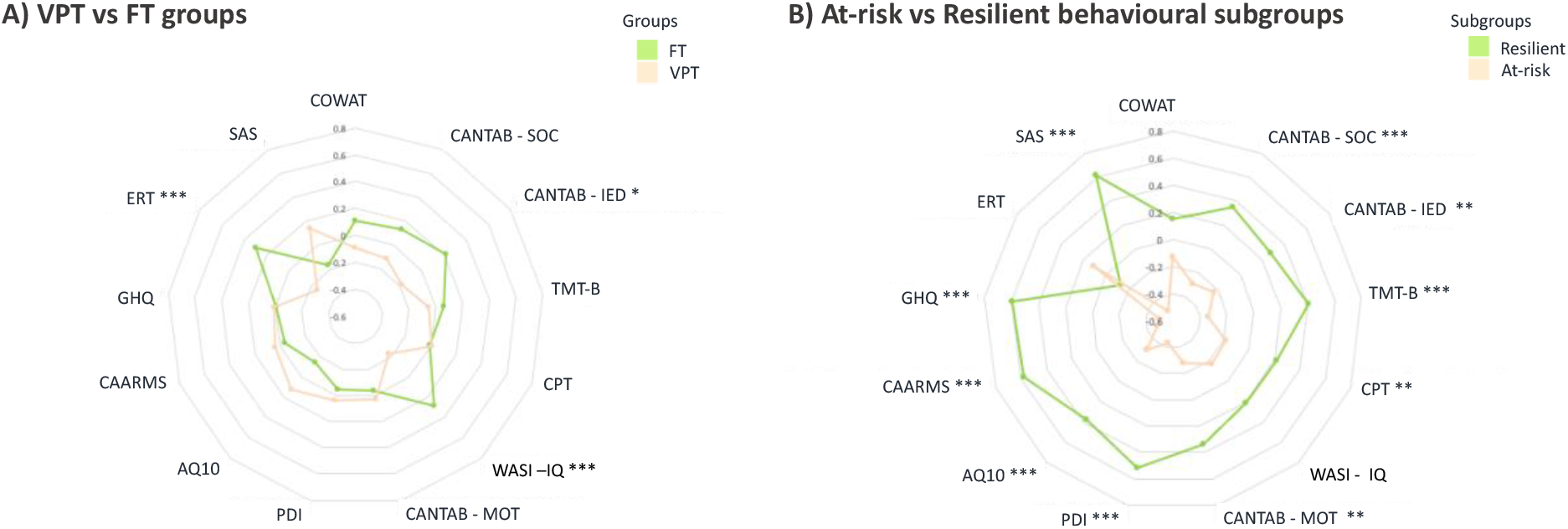
Radar plots showing differences in behavioural profiles between A) VPT and FT adults and B) At-risk and Resilient data-driven behavioural subgroups. Z-scores were computed for each group and plotted accordingly. For visual illustrative purposes, values for scales indicating poorer outcomes were reversed, so that larger Z-scores here indicate generally more optimal outcomes. *=p<0.05; **=p<0.01; ***p<0.001. Refer to Table 2 legend for behavioural measure abbreviations and Table SM 1 for descriptions.

**Table 1.** Clinical and socio-demographic characteristics of study participants.

|  | **VPT (n=116)** | **FT (n=83)** | **p-value** |
| --- | --- | --- | --- |
| **Gestational age at birth**, median (range) weeks | 30.00 (24.00 – 32.00) | n/a | n/a |
| **Birth weight**, median (range) grams | 1345 (552 – 2390) | 3440 (2690 – 4990) | < 0.001 |
| **Sex**, n (%) |  |  | 0.082 |
| *Male* | 66 (56.90%) | 36 (43.37%) |  |
| *Female* | 50 (43.10%) | 47 (56.63%) |  |
| <sup>a</sup> **Ethnicity**, n (%) |  |  | 0.139 |
| *African* | 2 (1.72%) | 5 (6.02%) |  |
| *Afro-Caribbean* | 2 (1.72%) | 4 (4.82%) |  |
| *Caucasian/White* | 76 (65.52%) | 55 (66.27%) |  |
| *Indian Subcontinent* | 8 (6.90%) | 2 (2.41%) |  |
| *Other* | 4 (3.45%) | 6 (7.23%) |  |
| <sup>b</sup> **Perinatal brain injury**, n (%) |  |  | n/a |
| *No injury* | 62 (53.45%) | n/a |  |
| *Minor injury* | 27 (23.28%) | n/a |  |
| *Major injury* | 26 (22.41%) | n/a |  |
| <sup>c</sup> **Parental socio-economic status at birth**, n (%) |  |  | 0.106 |
| *I – II* | 43 (51.81%) | 38 (32.76%) |  |
| *III* | 36 (43.37%) | 15 (12.93%) |  |
| *IV – V* | 8 (9.63%) | 3 (2.59%) |  |
| <sup>c</sup> **Participants' current socio-economic status, n (%)** |  |  | < 0.001 |
| *I – II* | 51 (43.97%) | 36 (43.37%) |  |
| *III* | 41 (35.35%) | 26 (31.33%) |  |
| *IV – V* | 6 (5.17%) | 0 (0.00%) |  |
| *Student* | 1 (0.86%) | 16 (19.28%) |  |
| *Unemployed* | 16 (13.8%) | 4 (4.82%) |  |
| **Age at assessment, median (range) years** | 31.37 (23.346 – 39.33) | 28.73 (26.26 – 36.49) | < 0.001 |

**Note.** <sup>a</sup> Ethnicity was self-reported. <sup>b</sup> Ultrasound scans were used to classify perinatal brain injury into three categories: no haemorrhage (no injury), grade I – II periventricular haemorrhage without ventricular dilation (minor injury) and grade III – IV periventricular haemorrhage with ventricular dilation (major injury). <sup>c</sup> Socio-economic status was categorised according to the Office of National Statistics, 1980 occupation classifications. I: Higher managerial, administrative and professional occupations; II: Intermediate occupations, small employers and own account workers; III: Routine and manual occupations – lower supervisory and technical and semi-routine and routine occupations. Missing data: 1 VPT and 1 FT had missing socio-economic status data; 24 VPT and 11 FT had missing ethnicity data; 1 VPT has missing perinatal brain injury classification.

**Table 2.** Behavioural outcomes in VPT and FT adults.

|  | VPT (n=116) | FT (n=83) | p-value | FDR p-value | Adj. FDR p-value | Effect size |
| --- | --- | --- | --- | --- | --- | --- |
| *COWAT, total words* | 13.00 (5.75) | 14.00 (5.25) | 0.052 | 0.166 | 0.115 | -0.042 |
| *CANTAB – SOC, problems solved* | 9.00 (2.75) | 10.00 (2.00) | 0.063 | 0.166 | 0.106 | -0.064 |
| *CANTAB – IED, total errors adjusted* | 15.00 (25.50) | 10.50 (14.65) | 0.002 | 0.007 | 0.021 | 0.184 |
| *TMT-B, time to finish task* | 73.50 (40.50) | 71.30 (39.05) | 0.081 | 0.175 | 0.068 | -0.093 |
| *CPT, total reaction time for correct responses* | 417.50 (59.15) | 414.00 (54.40) | 0.921 | 0.921 | 0.936 | -0.009 |
| *WASI – full scale IQ* | 106.00(13.75) | 113.50 (12.25) | <0.001 | <0.001 | <0.001 | 0.088 |
| *CANTAB – MOT, reaction time* | 691.00 (200.80) | 734.00 (196.90) | 0.307 | 0.399 | 0.456 | 0.062 |
| *PDI, total score* | 21.50 (50.25) | 18.00 (39.25) | 0.406 | 0.480 | 0.530 | 0.002 |
| *AQ10, total score* | 2.00 (2.44) | 3.00 (2.32) | 0.198 | 0.322 | 0.257 | 0.121 |
| *CAARMS, general psychopathology score* | 2.00 (5.50) | 2.00 (4.00) | 0.232 | 0.335 | 0.220 | -0.111 |
| *GHQ, total score* | 10.00 (6.00) | 10.00 (7.00) | 0.891 | 0.921 | 0.943 | 0.070 |
| *ERT, total correct* | 56.60 (11.15) | 62.00 (9.45) | <0.001 | <0.001 | <0.001 | 0.358 |
| *SAS, total score* | 1.58 (0.45) | 1.69 (0.53) | 0.127 | 0.236 | 0.4021 | 0.136 |

**Note.** Median (interquartile range) reported. “Adj. FDR p-value” corresponds to the p-value after adjusting for covariates (sex, age, socio-economic status) and correcting for multiple comparisons with FDR. Effect sizes are calculated using Wilcoxon Glass Rank Biserial Correlation. Missing data: <sup>a</sup> FT n=7, VPT n=22; <sup>b</sup> FT n=21, VPT n=22; <sup>c</sup> FT n=21, VPT n=19; <sup>d</sup> FT n=12, VPT n=17; <sup>e</sup> FT n=5, VPT n=9. Abbreviations. AQ10 = Autism Quotient; CANTAB = Cambridge Neurophysiological Test Automated Battery; CAARMS = Comprehensive Assessment of At-Risk Mental States; COWAT = Controlled Oral Word Association Test; CPT = Continuous Performance Test; ERT = Emotion Recognition Task; FT= full-term; GHQ = General Health Questionnaire; IED = Intra-Extra Dimensional Set Shift; MOT = Motor Screening Task; PDI = Peters Delusion Inventory; SAS =

### 1.3.2 Data-driven behavioural subgroups

Two data-driven behavioural subgroups were identified and labelled as ‘At-risk’ and ‘Resilient’, based on their observed phenotypic profiles (Table 3; Figure 2B).

**Table 3.**
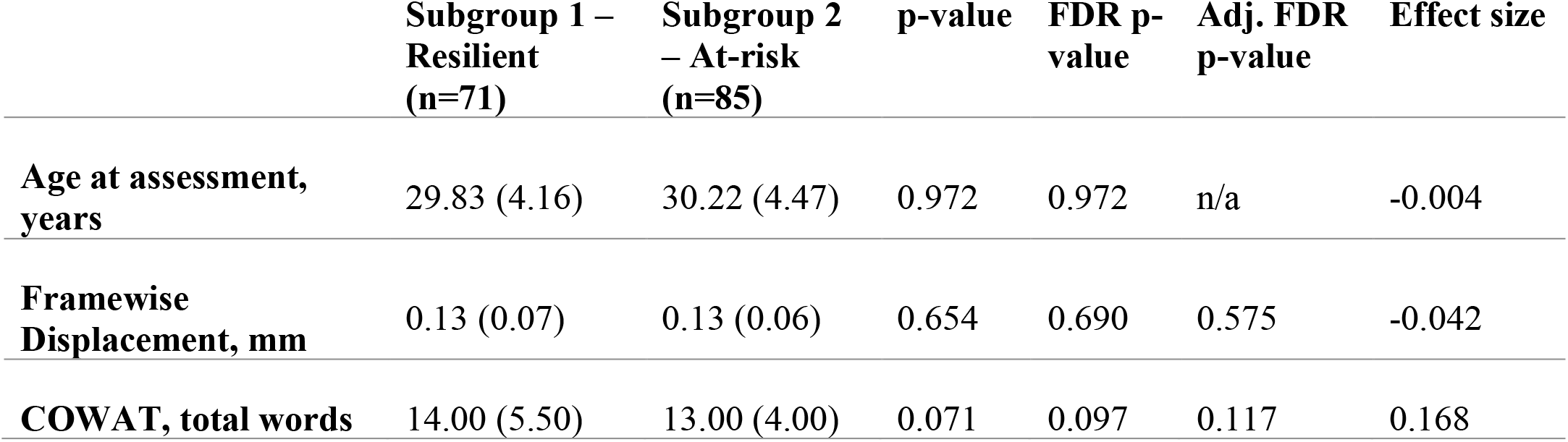
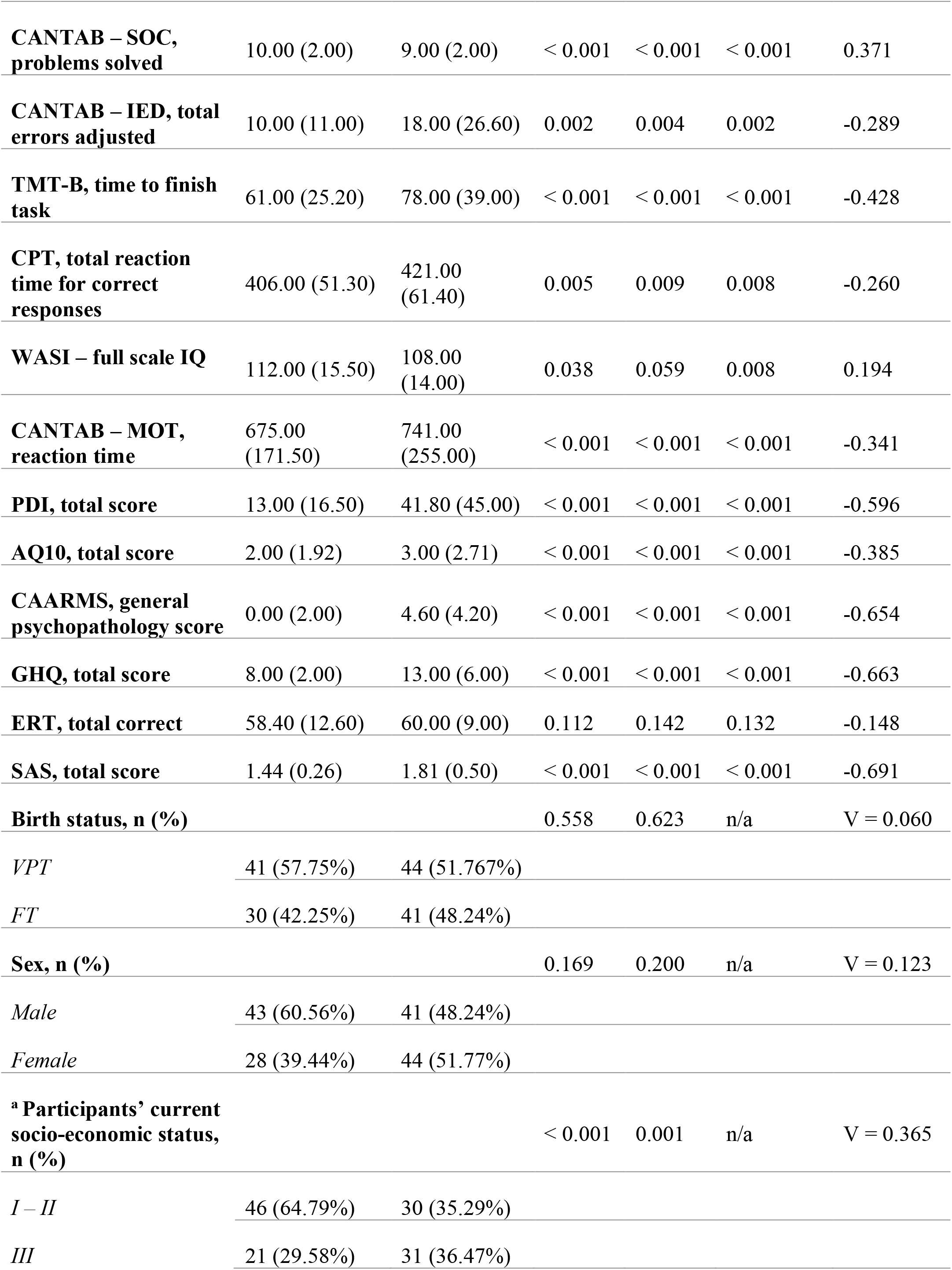
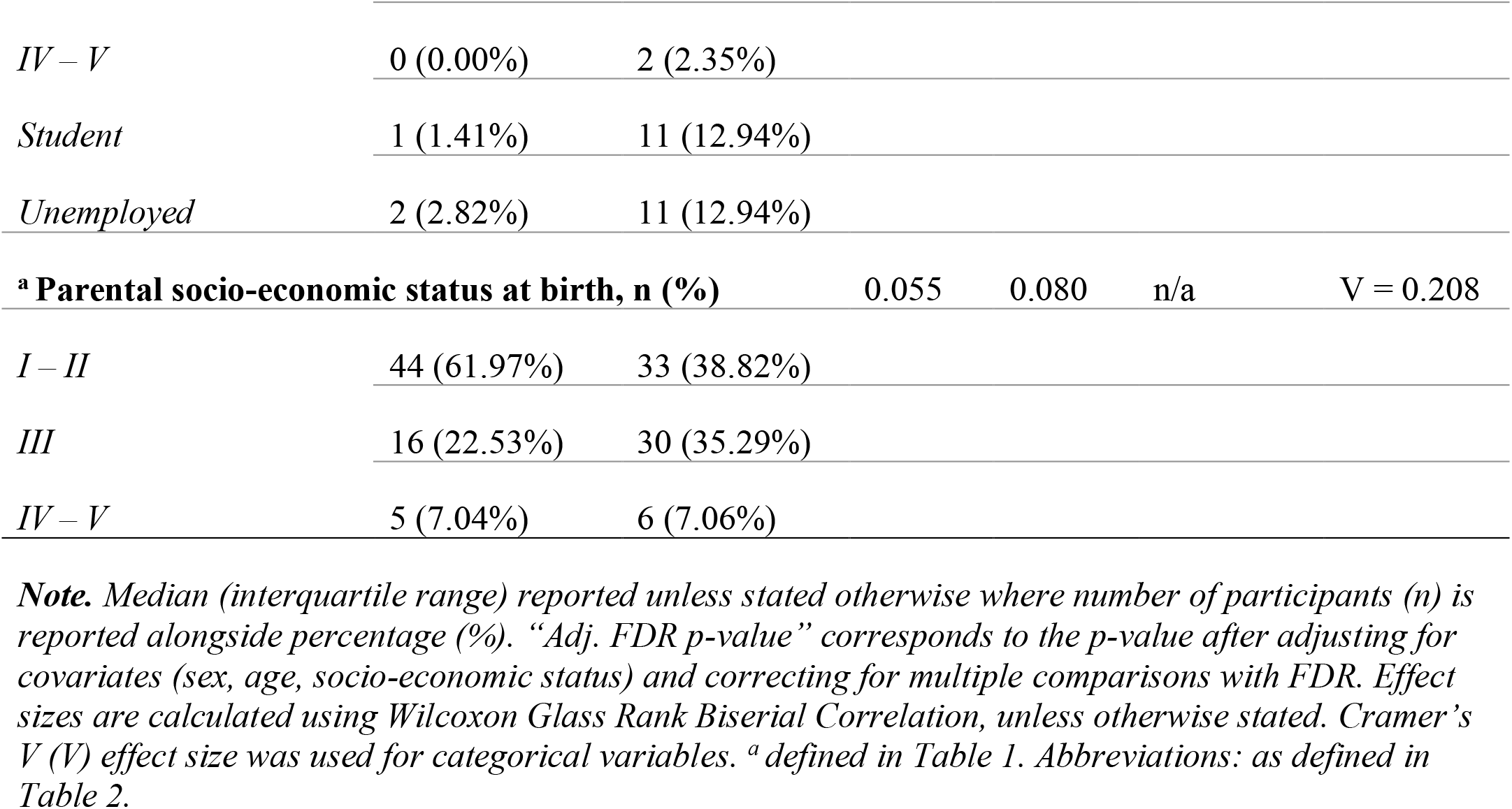
At-risk and Resilient behavioural subgroup profiles.

In summary, the At-risk subgroup had less optimal executive function and attention scores probing spatial planning, attentional set-shifting, visuo-motor coordination, comprehension abilities, sustained attention and response inhibition (CANTAB – SOC, MOT and IED, the TMT-B, and CPT), compared to the Resilient subgroup. The At-risk subgroup also had less optimal social adjustment, mental wellbeing, and psychiatric scores (PDI, CAARMS, GHQ, and SAS), and increased autistic traits (AQ-10 scores), compared to the Resilient subgroup. The two subgroups showed no differences in full-scale IQ (WASI), emotion recognition (ERT), or phonemic verbal fluency (COWAT). However, the At-risk subgroup had a higher proportion of individuals with lower own socio-economic status compared to the Resilient subgroup. Parental socio-economic status did not differ between the subgroups.

52% of the VPT adults in our sample clustered into the At-risk subgroup, and the remaining 48% into the Resilient subgroup (Figure 3). Upon examining VPT adults only, there were no significant differences between the At-risk and Resilient subgroups in terms of perinatal clinical measures (i.e., gestational age, birth weight, or perinatal brain injury) (Table SM 3; Figure SM 3). In terms of parental socio-economic status, there were no differences between At-risk and Resilient subgroups within VPT or FT adults (Table SM 3 and Table SM 4, respectively). As for participants’ own socio-economic status, only those born VPT displayed differences between the data-driven behavioural subgroups, where more VPT individuals with higher managerial, administrative, and professional occupations belonged the Resilient subgroup compared to the At-Risk subgroup (Table SM 3). However, socio-economic status for those born FT did not differ significantly between the two data-driven subgroups (Table SM 4).

**Figure 3.**
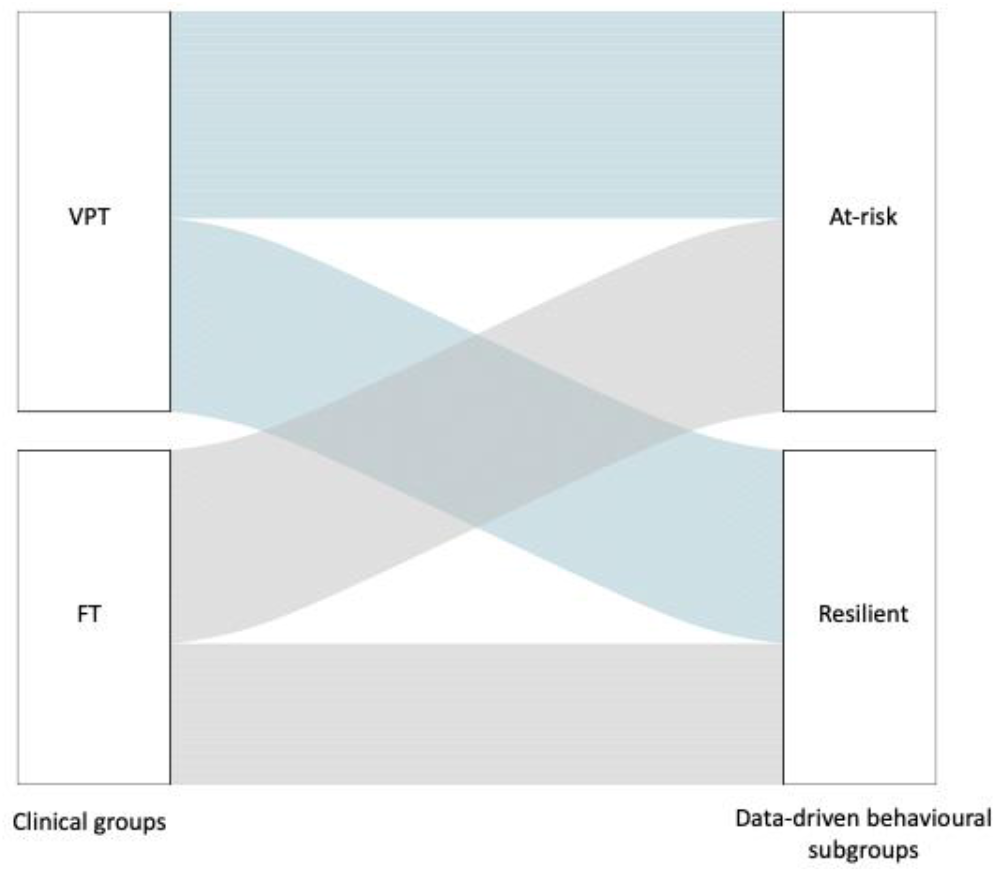
Alluvial plot showing VPT (in blue) and FT (in grey) adults clustering into the At-risk and Resilient data-driven behavioural subgroups.

### 1.3.3 Between-group differences in rsFC

We report NBS analyses using p-NBS-Threshold values powered to detect a significant effect, whilst also reducing component size (i.e., not p = 0.05) (Table SM 5). Main results reported here are from one-tailed NBS analyses using p-NBS-Threshold = 0.01, and additional sensitivity analyses investigating rsFC using a more stringent threshold (p-NBS-Threshold = 0.001) are reported in Supplementary Material.

#### VPT < FT

NBS results showed weaker rsFC in the VPT group compared to the FT group (i.e., VPT < FT) in one component comprising 360 nodes (i.e., 96.25% of all regions) and 1467 edges (i.e., 2.10% of the 69,751 possible connections), with a component strength of 616.04 (p-FWER value = 0.007). Regions included in this component were widespread across the brain (Figure 4A; Table SM 6). Nodes with the highest number of connections within the component (i.e., component ‘hub’ regions) were predominantly localised to superior temporal gyrus, inferior and superior parietal cortex, inferior frontal, orbitofrontal, anterior cingulate and medial prefrontal cortex, inferior premotor, a lateral occipital/posterior temporal visual area, dorsolateral prefrontal cortex, medial and lateral temporal, and posterior cingulate cortex. Component within- and between-network connectivity was highest in the DMN (Figure 5A).

**Figure 4.**
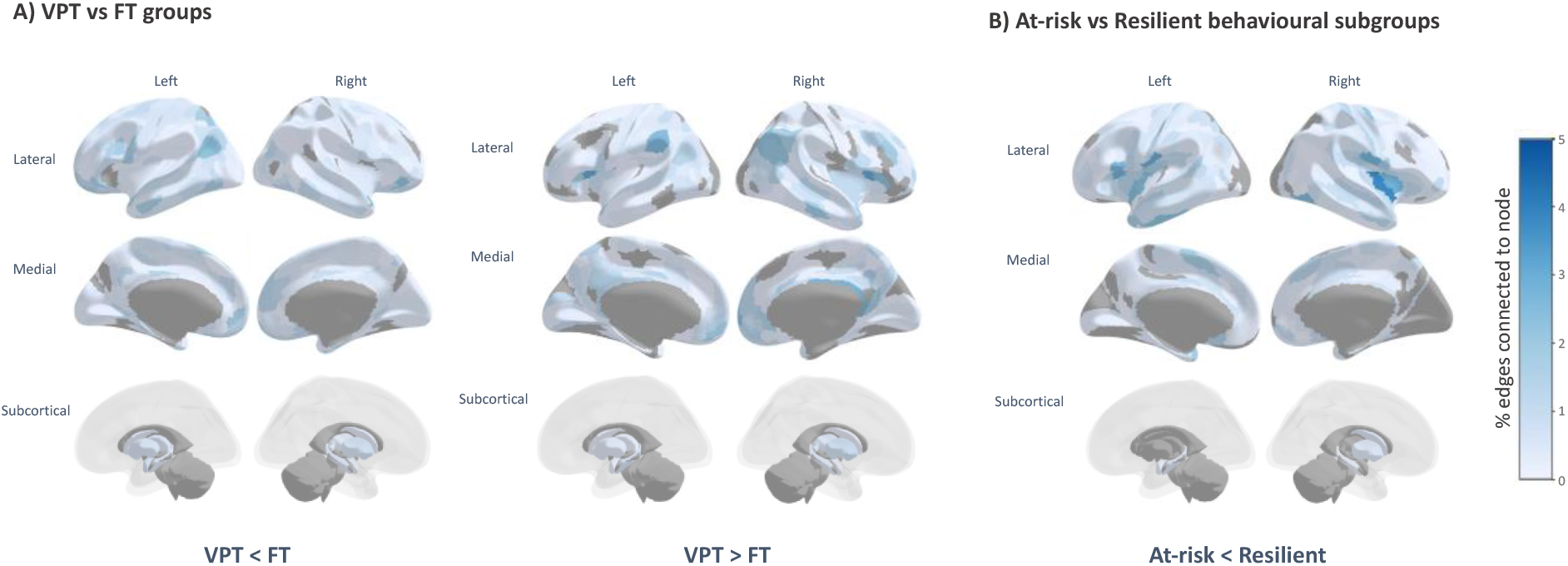
Percentage of edges connected to each region (i.e., node) within the significant NBS components for A) VPT vs FT groups and B) At-risk vs Resilient behavioural subgroups. Main analysis results from NBS modelling using a p-NBS-Threshold of 0.01, 1000 permutations, and linear models correcting for covariates (age, sex, in-scanner head motion, and socio-economic status). Darker colours (blue) denote higher percentages of edges and lighter colours (light blue and white) denote lower percentages, with areas marked in grey indicating regions that are not forming part of the NBS component.

**Figure 5.**
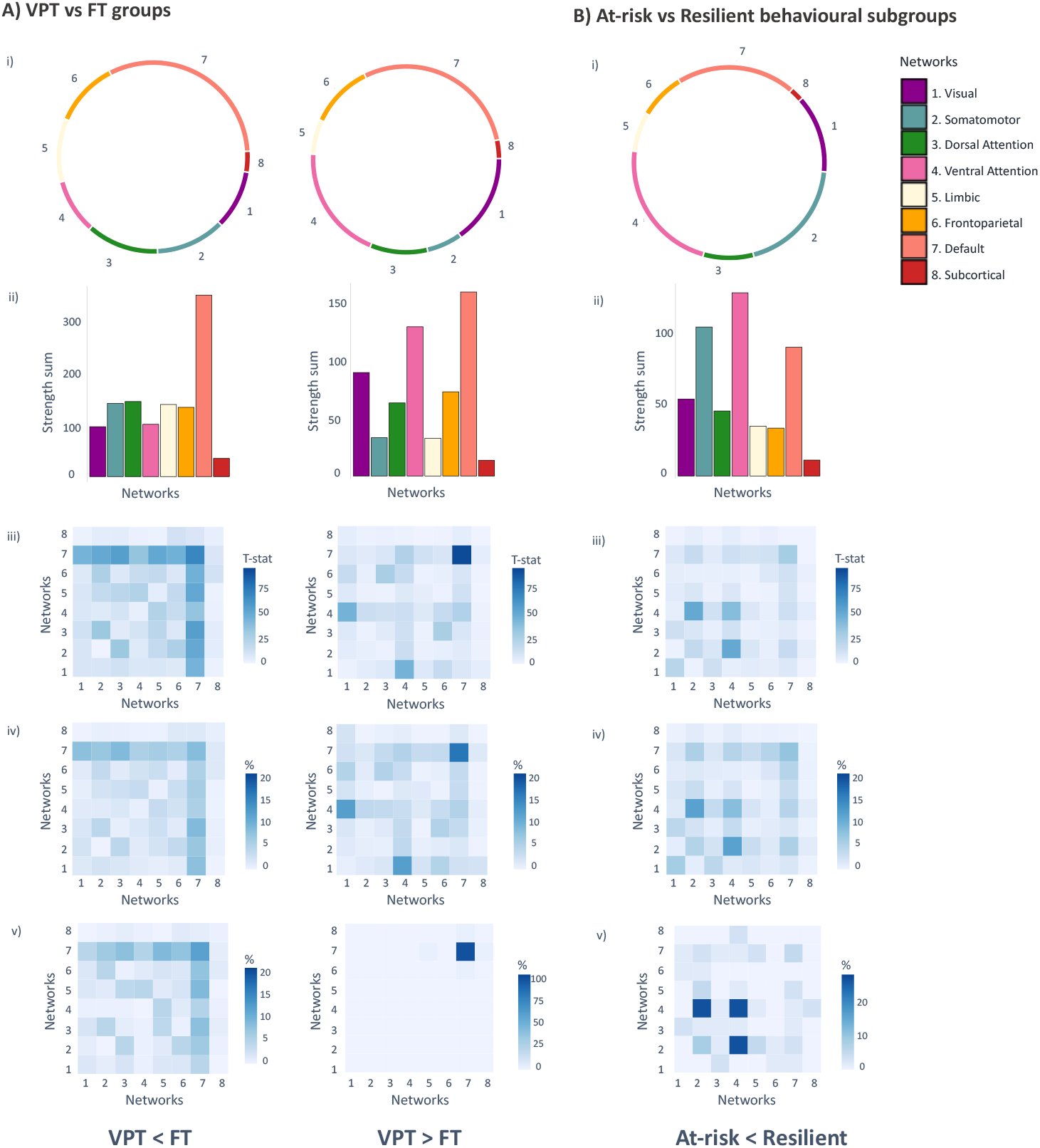
Within- and between-network connectivity of the significant NBS components in A) VPT vs FT groups and B) At-risk vs Resilient behavioural subgroups. Results from main NBS analyses using a p-NBS-threshold of 0.01: i) circle plots illustrating within- and between-network connections within the significant component only; ii) bar plots showing the sum of T-statistic strength values within the significant NBS component belonging to the different intrinsic connectivity networks (i.e., seven Yeo networks and an eighth network of subcortical regions), and iii) within- and between-network connectivity strength (T-statistic sum). Heatmaps showing total number of within- and between-network connections as a percentage of the total number of connections forming the significant component: iv) at p-NBS-threshold = 0.01, and v) p-NBS-threshold = 0.001.

#### VPT > FT

NBS results also showed greater rsFC in the VPT group compared to the FT group (i.e., VPT > FT) in one component comprising 340 nodes (i.e., 90.91% of regions), 962 edges (i.e., 1.37% of possible connections) and component strength of 358.03 (p-FWER value < 0.001). ‘Hub’ regions within this component were less widespread across the brain and localised within posterior opercular cortex, posterior cingulate cortex, inferior parietal cortex, right orbitofrontal cortex, bilateral anterior cingulate and medial prefrontal cortex, superior temporal gyrus (auditory association cortex), dorsolateral prefrontal cortex, right lateral temporal cortex, right temporo-parietal-occipital junction, and medial superior parietal cortex (Figure 4A; Table SM 7). The highest number of connections found in the component were within the DMN itself, followed by a moderate number of widespread connections in the VAN, and especially between the VAN and the visual network.

A total of 326 nodes (i.e., 87.17% of regions) were present in both VPT < FT and VPT > FT components; however, the sets of edges connecting nodes within each component were mutually exclusive with no overlapping edges.

#### At-Risk < Resilient

Contrasts testing for lower rsFC in the At-risk compared to the Resilient subgroup identified one significant NBS component with 337 nodes (i.e., 90.11% of regions), 832 edges (i.e., 1.19% of possible connections) and a strength sum of 309.04 (p-FWER = 0.019). Hub regions with the highest number of connections within the component were predominantly located in insular, frontal opercular, and posterior opercular cortex (Figure 4B; Table SM 8). Other hub regions were found in the left inferior frontal cortex, lateral temporal cortex, right temporo-occipital visual area, left temporo-parieto-occipital junction, anterior cingulate, medial prefrontal cortex, left supplementary motor area, primary somatosensory cortex, and the superior temporal sulcus (auditory association cortex) (Figure 4B; Table SM 8). Component within- and between-network connectivity were most pronounced between the VAN and somatomotor networks, and within the VAN (Figure 5B).

#### At-Risk > Resilient

No significant NBS components were detected when testing for higher rsFC in the At-risk compared to the Resilient subgroup.

Confirming the robustness of the observed effects from analyses using a p-NBS-threshold of 0.01, sensitivity NBS analyses using a more stringent p-NBS-threshold of 0.001 reported significant components with greater sparsity (Table SM 9), but largely similar rsFC patterns (Figure 4; Figure 5Av; Figure 5Bv).

Post-hoc exploratory analyses investigating the interaction between clinical (VPT vs FT) groups and data-driven behavioural subgroups (At-risk vs Resilient) on rsFC did not identify significant components (p-FWER > 0.05) at any p-NBS-Threshold examined (0.05, 0.01, and 0.001). Similarity index calculations indicated that the At-risk < Resilient component had a high number of nodes, which were also part of the VPT < FT component (n=325; Sørensen-Dice = 0.93) and the VPT > FT component (n=304; Sørensen-Dice = 0.90), but very few edges overlapped with either clinical component; n=9 edges (Sørensen-Dice = 0.01) and n=22 edges (Sørensen-Dice = 0.03), respectively.

## 1.4 Discussion

In this study, we compared rsFC between groups of adults stratified in terms of (i) their clinical characteristics (i.e., VPT and FT birth) as well as (ii) their behavioural profiles identified using data-driven consensus clustering, regardless of their gestational age at birth. In VPT compared to FT adults, we identified complex preterm-specific patterns of both increased and decreased intrinsic rsFC predominately characterised by hypo-connectivity between the DMN and other networks examined and hyper-connectivity within the DMN and between the VAN and the visual network. When VPT and FT born adults were stratified in terms of their data-driven behavioural profiles, irrespective of gestational age at birth, we identified an ‘At-risk’ subgroup with more behavioural difficulties and reduced rsFC anchored in frontal opercular and insular areas of the VAN, relative to a ‘Resilient’ subgroup with more favourable behavioural outcomes.

In summary, our results indicate that there are complex and widespread long-lasting preterm-specific rsFC alterations, which we speculate may confer both risk and resilience to the behavioural sequelae associated with VPT birth. That is, while these rsFC alterations may partly explain the behavioural difficulties specific to those born VPT in cognitive and socio-emotional processing observed here, they may also aid the preservation of optimal outcomes in other behavioural domains where no between-group differences were noted (e.g., psychiatric difficulties, sustained attention, planning or phonemic verbal fluency). On the other hand, localised functional hypo-connectivity anchored in insular and frontal opercular regions observed in our study may characterise participants with unfavourable compared to favourable cognitive and behavioural outcomes, irrespective of birth status.

### 1.4.1 Differences in rsFC and behavioural outcomes between VPT and FT born adults

We identified complex patterns of both hypo- and hyper-connectivity predominantly located in the DMN, VAN, and visual networks in VPT compared to FT participants. Such rsFC alterations are evident in adulthood and may represent the neurobiological architecture underlying the attentional, cognitive, and socio-emotional processing difficulties associated with VPT birth, commonly referred to as the “preterm behavioural phenotype” (Johnson and Marlow, 2011). However, in our cohort, VPT relative to FT born adults only differed in selected dimensions that have been studied as part of the “preterm behavioural phenotype”; they had lower full-scale IQ, difficulties in rule learning, attentional set shifting abilities (measured by the CANTAB IED), and emotion recognition.

VPT adults, compared to controls, displayed functional hypo-connectivity between the DMN and the visual, somatomotor, dorsal attention, limbic and frontoparietal networks, as well as hyper-connectivity within the DMN itself. In line with our findings, patterns of both hyper- and hypo-connectivity in the DMN have been previously reported in VPT born children and adults (Bäuml et al., 2015; Degnan et al., 2015; Mossad et al., 2022; Wheelock et al., 2021), suggesting that functional DMN connectivity alterations may be characteristic of VPT samples. Functional DMN connectivity emerges during the third trimester of gestation, a critical period of brain development during which VPT infants are born, and previous studies have reported structural and functional brain alterations at term-equivalent age in regions belonging to the DMN (Doria et al., 2010; Sa de Almeida et al., 2021; Scheinost et al., 2022; Smyser et al., 2016, 2010). Extending beyond preterm populations, functional alterations in the DMN have been described in several psychiatric conditions, including schizophrenia, anxiety, and mood disorders (Buckner, 2013; Doucet et al., 2020), suggesting that the DMN rsFC alterations observed in VPT individuals may represent neurobiological changes which could contribute to the behavioural difficulties associated with VPT birth.

On the other hand, alterations in DMN rsFC have also been studied as adaptive neural mechanisms; for instance, maintaining attentional capture (i.e., less distractibility) in male veterans (Poole et al., 2016). Such findings suggest that functional reorganisation of the DMN may also reflect compensatory biological alterations supporting selective cognitive and behavioural processing in VPT individuals; in this context referring to the behavioural outcomes where no between-group differences were noted in our study sample, including spatial planning (CANTAB – SOC), coordination (MOT), cognitive flexibility (TMT-B), phonemic verbal fluency (COWAT), sustained attention (CPT), social adaptation (SAS), prodromal symptoms (PDI), autism traits (AQ10) and general psychopathology (CAARMS and GHQ). This finding emphasises the notion that complex neurobiological alterations following VPT birth may confer both risk and resilience to the long-term consequences of VPT birth. Further supporting this point, we also identified patterns of hyper-connectivity in the VPT relative to the FT group in the VAN, a “circuit-breaker” network which disengages during tasks requiring focused attention and activates to redirect attention towards external task-irrelevant stimuli (Corbetta and Shulman, 2002; Vossel et al., 2014). Notably, the highest proportion of connections were between the VAN and the visual network, which may reflect adaptive functional reorganisation in the VPT group. In a previous study, stronger rsFC changes in visual and attention networks were associated with fewer attention deficits in visual short-term memory storage in VPT relative to FT adults (Finke et al., 2015). Another study found that attention processing was selectively supported by VAN and visual network connectivity in VPT born children, and by dorsal attention, frontoparietal, and cingulo-opercular network connectivity in FT controls (Wheelock et al., 2021). The authors argued that VPT children may have a greater reliance on visually stimulated “bottom-up” neural processes to maintain attention mechanisms, which is in line with their previous findings showing poorer attention abilities in VPT children with reduced volumes in regions of the visual network (Lean et al., 2017).

We also identified that component ‘hub’ regions (i.e., those with a high percentage of connections within the component) with higher rsFC in the VPT group relative to the FT group, were localised to brain regions previously identified as nodes of a ‘rich-club’ network (i.e., the sub-network of highly connected brain regions which are also highly connected to one another), important for efficient integration and transfer of information between systems (Grayson et al., 2014; van den Heuvel and Sporns, 2013). We previously reported stronger rich-club network structural connectivity and weaker peripheral connectivity in an overlapping sample of VPT adults compared to FT controls, and argued that increased resources in the VPT brain may be preferentially allocated to the rich-club network in order to maintain efficient information exchange across the brain (Karolis et al., 2016).

### 1.4.2 Differences in rsFC and behavioural outcomes between data-driven behavioural subgroups

Considering the neurodevelopmental heterogeneity exhibited within and between those born VPT and FT, it remains to be established whether we can use rsFC to characterise the behavioural difficulties observed in VPT individuals (Anderson et al., 2021; Nosarti et al., 2012). Aiming to address this question, we stratified VPT and FT adults into data-driven behavioural subgroups and investigated specific rsFC alterations which may differentiate between them. We identified two data-driven behavioural subgroups, irrespective of birth status (VPT and FT): an ‘At-risk’ subgroup with more executive function, attention, socio-emotional, and psychiatric difficulties, compared to a ‘Resilient’ subgroup, with more favourable behavioural outcomes. Notably, the behavioural differences observed between data-driven subgroups were more pronounced than those observed between VPT and FT adults.

We also identified underlying rsFC differences characterising the distinct data-driven behavioural subgroups, where the At-risk, compared to the Resilient subgroup, displayed hypo-connectivity within the VAN and between the VAN and the somatomotor network. Specifically, the predominant connectivity patterns forming this component were anchored in frontal opercular and insular regions of the brain, which play an integral role in detecting bottom-up salient information from the environment and switching between networks to produce appropriate cognitive control, socio-emotional, and interoceptive somatomotor responses (Deen et al., 2011; Higo et al., 2011; Loitfelder et al., 2015; Menon and Uddin, 2010; Quirmbach and Limanowski, 2022; Uddin et al., 2017). Our findings here are in line with previous studies showing structural and functional alterations in insular and opercular regions in adults experiencing mental health difficulties (Horn et al., 2010; Yin et al., 2018) and executive dysfunction (Hausman et al., 2022). Furthermore, studies investigating rsFC across multiple psychiatric groups identified transdiagnostic patterns of hypo-connectivity in lower-order networks, such as the somatomotor network, as well as higher order networks, such as the VAN (Baker et al., 2019; Li et al., 2021). The rsFC patterns identified here characterised data-driven behavioural subgroups irrespective of gestational age at birth (VPT and FT), indicating that these specific neural mechanisms may represent biomarkers of behavioural outcomes in the general population which are not unique to VPT individuals. We also found no significant interaction effects between birth group (VPT vs FT) and data-driven behavioural subgroups (At-risk vs Resilient) on rsFC and very little overlap in rsFC between the clinical and behavioural components identified by NBS, which may further support our speculation that the differences in rsFC between the data-driven subgroups may be characterising behavioural outcomes independently of gestational age at birth. However, future studies with larger samples, and hence greater statistical power, may further investigate the possible influence of VPT (vs FT) birth on the relationship between rsFC alterations and behavioural outcomes.

Our post-hoc analyses aimed to explore whether specific enriching factors, or lack of certain social or clinical risk factors, protected the VPT adults belonging to the Resilient subgroup from developing an At-risk behavioural profile. In contrast to previous studies in VPT children, we found that perinatal clinical risk was not higher in VPT adults who belonged to an At-risk (vs Resilient) subgroup (Hadaya et al., 2023; Poehlmann et al., 2015). Social risk, on the other hand, may be specifically related to the difficulties observed in the VPT At-risk subgroup, which contained more VPT adults from more socially disadvantaged backgrounds compared to the Resilient subgroup, while this relationship was not observed in FT adults. These findings as well as previous studies in children (Hadaya et al., 2023; Lean et al., 2020; Ross et al., 2016; Vanes et al., 2021) could be interpreted within a “differential susceptibility” framework, which posits that vulnerable individuals (e.g., those born VPT) are particularly sensitive to environmental influences, where negative or positive factors (such as social (dis)advantage) can promote either worse or more optimal outcomes, respectively (Belsky et al., 2007). Therefore, VPT adults in the At-risk subgroup may have experienced a “double-hit” of being born VPT as well as being socio-economically disadvantaged. Nonetheless, it is worth noting, that socio-economic status in our sample only partially explained behavioural outcomes, as our main behavioural and rsFC results remained significant after adjusting for this covariate. It is therefore plausible that additional unmeasured environmental or hereditary factors (e.g., parental mental health or cognitively stimulating home environment) (Hadaya et al., 2023; Lean et al., 2020; Vanes et al., 2021) may have contributed to the behavioural outcomes observed in the distinct subgroups.

This study has several strengths, which include the use of a large sample of both VPT and FT born controls, the implementation of rigorous consensus clustering methods to obtain data-driven behavioural subgroups, as well as the use of fMRIPrep, a robust automated resting state functional MRI pre-processing pipeline which promotes pre-processing transparency and aims to alleviate hurdles related to reproducibility in functional MRI analyses (Esteban et al., 2019; Pernet and Poline, 2015). We also acknowledge several limitations to our study. After excluding participants with excessive head motion, behavioural outliers, missing data, or poor alignment of functional MRI data, supplementary analyses showed that the subsample of VPT adults used in our analyses had relatively better cognitive and socio-emotional processing outcomes in comparison to VPT adults excluded from the analyses. This may limit the generalisability of our results to cohorts of low-risk VPT adults with relatively favourable behavioural outcomes. It may also explain why our two data-driven behavioural subgroups have similar proportions of VPT and FT born individuals, which is not in line with previous studies in children that have reported higher ratios of VPT to FT individuals belonging to At-risk subgroups and lower ratios to Resilient subgroups (Burnett et al., 2019; Lean et al., 2020). On the other hand, our results may be reflective of the increased rates of mental health difficulties with increasing age, which may not yet be apparent in childhood (Otto et al., 2021; Solmi et al., 2022). Future studies with more representative samples of VPT adults could help elucidate these potentially inconsistent findings. Another possible limitation is that we did not include known risk factors (such as socio-economic status, parenting or clinical measures) in the clustering model, which may have increased the difficulty in identifying nuanced subgroups exhibiting ‘equifinal’ trajectories (i.e., those with similar behavioural outcomes but distinct underlying risk factors) (Cicchetti and Rogosch, 1996; Hadaya et al., 2023). However, to our knowledge, this is the first study to parse behavioural heterogeneity in VPT adults; therefore, we decided to follow an approach similar to those implemented in the vast majority of studies in VPT children, where individual-level behavioural variables were included as inputs to the clustering model and risk factors were explored post-hoc (Bogičević et al., 2021; Burnett et al., 2019; Johnson et al., 2018; Lean et al., 2020; Poehlmann et al., 2015; Ross et al., 2016; van Houdt et al., 2020).

In summary, this study shows that there are complex patterns of rsFC alterations which are specifically associated with VPT birth in adult life. We speculate that these alterations may reflect neural adaptations conferring both risk and resilience to the long-term sequelae of VPT birth. We also identify distinct rsFC alterations in insular and frontal opercular regions in a data-driven At-risk relative to a Resilient behavioural subgroup, irrespective of birth status (VPT vs FT), indicating that these neurobiological changes may reflect biomarkers of behavioural outcomes in the general population that are not unique to those born VPT.

## Acknowledgements

The authors would like to thank the participants and their families for taking part in the research study, and the clinical, research and radiography teams for help with data collection. The authors acknowledge the use of the Computational Research, Engineering and Technology Environment (CREATE) research computing facility at King’s College London.(King*’*s College London, 2022)

## Funding

The work was supported by the Medical Research Council (MRC), UK [grant number: MR/K004867/1]. The authors acknowledge infrastructure support from the National Institute for Health Research (NIHR) Mental Health Biomedical Research Centre (BRC) at South London, Maudsley NHS Foundation Trust and Institute of Psychiatry, Psychology and Neuroscience, King’s College London. The authors also acknowledge support in part from the Engineering and Physical Sciences Research Council (EPSRC) Centre for Medical Engineering at Kings College London [WT 203148/Z/16/Z], MRC strategic grant [MR/K006355/1], the Department of Health through an NIHR Comprehensive Biomedical Research Centre Award (to King’s College Hospital NHS Foundation Trust).

## Competing interests

Authors declare no potential conflicts of interest.

## Data availability

Access to data supporting the published work can be made available upon request from the corresponding author. Code used to label nodes according to intrinsic network membership is accessible here: https://github.com/frantisekvasa/functional_network_development/blob/master/nspn.fmri.R and code used to run consensus clustering pipelines is adapted from code accessible here: https://github.com/lailahadaya/preterm-ExecuteSNF.CC.

## Notes

### Competing Interest Statement

The authors have declared no competing interest.

