## Supplementary Material document for "Exploring functional connectivity in clinical and data-driven groups of preterm and term adults"

**Supplementary Material document for manuscript entitled**: Exploring functional connectivity in clinical and data-driven groups of preterm and term adults

**Running title:** Preterm and term adult brain-behaviour

**Authors**: Laila Hadaya ([1,2]), František Váša ([3]), Konstantina Dimitrakopoulou ([4]), Mansoor Saqi ([4]), Sukhwinder S Shergill ([5,6,7]), A David Edwards ([1]), Dafnis Batalle ([1,8]), Robert Leech ([3]), Chiara Nosarti ([1,2]).

**Affiliations**: [1]Centre for the Developing Brain, Department of Perinatal Imaging and Health, King's College London (KCL), London, United Kingdom. [2]Department of Child and Adolescent Psychiatry, Institute of Psychiatry Psychology and Neuroscience (IoPPN), KCL, London, United Kingdom. [3]Department of Neuroimaging, IoPPN, KCL, London, United Kingdom. [4]Translational Bioinformatics Platform, NIHR Biomedical Research Centre, Guy's and St. Thomas' NHS Foundation Trust and KCL, London, United Kingdom. [5]Kent and Medway Medical School, Canterbury, UK. [6]Kent and Medway NHS and Social Care Partnership Trust, Maidstone, UK. [7]Department of Psychosis Studies, IoPPN, KCL, London, United Kingdom. [8]Department of Forensic and Neurodevelopmental Sciences, IoPPN, KCL, London, United Kingdom.

**Correspondence to**: Chiara Nosarti, Centre for the Developing Brain, Department of Perinatal Imaging and Health, School of Biomedical Engineering & Imaging Sciences, King's College London, First Floor South Wing, St Thomas' Hospital, London SE1 7EH.

**Anatomical and functional MRI data pre-processing with fMRIPrep**

The full description of the anatomical and functional data pre-processing pipeline below is extracted from the boilerplate automatically generated by fMRIPrep (released under the CC0 license).

***Anatomical data pre-processing.*** The T1-weighted (T1w) image was corrected for intensity non-uniformity (INU) with N4BiasFieldCorrection (Tustison et al., 2010), distributed with ANTs 2.2.0 (RRID:SCR_004757, (Avants et al., 2008)), and used as T1w-reference throughout the workflow. The T1w-reference was then skull-stripped with a Nipype implementation of the antsBrainExtraction.sh workflow (from ANTs), using OASIS30ANTs as target template. Brain tissue segmentation of cerebrospinal fluid (CSF), white-matter (WM) and gray-matter (GM) was performed on the brain-extracted T1w using fast (FSL 5.0.9, RRID:SCR_002823, (Zhang et al., 2001)). Brain surfaces were reconstructed using recon-all (FreeSurfer 6.0.1, RRID:SCR_001847, (Dale et al., 1999)), and the brain mask estimated previously was refined with a custom variation of the method to reconcile ANTs-derived and FreeSurfer-derived segmentations of the cortical gray-matter of Mindboggle (RRID:SCR_002438, (Klein et al., 2017)). Volume-based spatial normalization to one standard space (MNI152NLin2009cAsym) was performed through nonlinear registration with antsRegistration (ANTs 2.2.0), using brain-extracted versions of both T1w reference and the T1w template. The following template was selected for spatial normalization: ICBM 152 Nonlinear Asymmetrical template version 2009c [(Fonov et al., 2011), RRID:SCR_008796; TemplateFlow ID: MNI152NLin2009cAsym].

***Functional data pre-processing.*** For each of the 1 BOLD runs found per subject (across all tasks and sessions), the following preprocessing was performed. First, a reference volume and its skull-stripped version were generated using a custom methodology of fMRIPrep. Head-motion parameters with respect to the BOLD reference (transformation matrices, and six corresponding rotation and translation parameters) are estimated before any spatiotemporal filtering using mcflirt (FSL 5.0.9, (Jenkinson et al., 2002). BOLD runs were slice-time corrected using 3dTshift from AFNI 20160207 (Cox and Hyde 1997, RRID:SCR_005927). Susceptibility distortion correction (SDC) was omitted. The BOLD reference was then co-registered to the T1w reference using bbregister (FreeSurfer) which implements boundary-based registration (Greve and Fischl, 2009). Co-registration was configured with six degrees of freedom. The BOLD time-series (including slice-timing correction when applied) were resampled onto their original, native space by applying the transforms to correct for head-motion. These resampled BOLD time-series will be referred to as preprocessed BOLD in original space, or just preprocessed BOLD. The BOLD time-series were resampled into standard space, generating a preprocessed BOLD run in MNI152NLin2009cAsym space. First, a reference volume and its skull-stripped version were generated using a custom methodology of fMRIPrep. Several confounding time-series were calculated based on the preprocessed BOLD: framewise displacement (FD), DVARS and three region-wise global signals. FD was computed using two formulations following Power (absolute sum of relative motions, (Power et al., 2014)) and Jenkinson (relative root mean square displacement between affines, (Jenkinson et al., 2002)). FD and DVARS are calculated for each functional run, both using their implementations in Nipype (following the definitions by (Power et al., 2014)). The three global signals are extracted within the CSF, the WM, and the whole-brain masks. Additionally, a set of physiological regressors were extracted to allow for component-based noise correction (CompCor, (Behzadi et al., 2007)). Principal components are estimated after high-pass filtering the preprocessed BOLD time-series (using a discrete cosine filter with 128s cut-off) for the two CompCor variants: temporal (tCompCor) and anatomical (aCompCor). tCompCor components are then calculated from the top 5% variable voxels within a mask covering the subcortical regions. This subcortical mask is obtained by heavily eroding the brain mask, which ensures it does not include cortical GM regions. For aCompCor, components are calculated within the intersection of the aforementioned mask and the union of CSF and WM masks calculated in T1w space, after their projection to the native space of each functional run (using the inverse BOLD-to-T1w transformation). Components are also calculated separately within the WM and CSF masks. For each CompCor decomposition, the k components with the largest singular values are retained, such that the retained components time series are sufficient to explain 50 percent of variance across the nuisance mask (CSF, WM, combined, or temporal). The remaining components are dropped from consideration. The head-motion estimates calculated in the correction step were also placed within the corresponding confounds file. The confound time series derived from head motion estimates and global signals were expanded with the inclusion of temporal derivatives and quadratic terms for each (Satterthwaite et al., 2013). Frames that exceeded a threshold of 0.5 mm FD or 1.5 standardised DVARS were annotated as motion outliers. All resamplings can be performed with a single interpolation step by composing all the pertinent transformations (i.e. head-motion transform matrices, susceptibility distortion correction when available, and coregistrations to anatomical and output spaces). Gridded (volumetric) resamplings were performed using antsApplyTransforms (ANTs), configured with Lanczos interpolation to minimize the smoothing effects of other kernels (Lanczos, 1964). Non-gridded (surface) resamplings were performed using mri_vol2surf (FreeSurfer).

Many internal operations of fMRIPrep use Nilearn 0.6.2 (Abraham et al., 2014), RRID:SCR_001362), mostly within the functional processing workflow. For more details of the pipeline, see the section corresponding to workflows in fMRIPrep's documentation (https://fmriprep.org/en/latest/workflows.html).

**Behavioural data pre-processing and consensus clustering feature selection**

Behavioural variables with data missing in >25% of the sample (i.e., total RFS scores) or with outlier data point values in >5% of the sample (i.e., PAL task total adjusted errors score and HSCT total scores) were excluded from the analyses. Participants were excluded from the analyses if they had more than 25% of behavioural data missing or had outlier data points on any of the included behavioural measures. Any remaining missing data were imputed using the K-nearest neighbour method. The final sample used in the consensus clustering pipeline excluded participants not included in the functional connectivity analysis and those with further outlier data points on any of the included behavioural measures (Figure SM 1). Outliers are defined as data points with values exceeding the median by three times the interquartile range or more.

### Supplementary figures

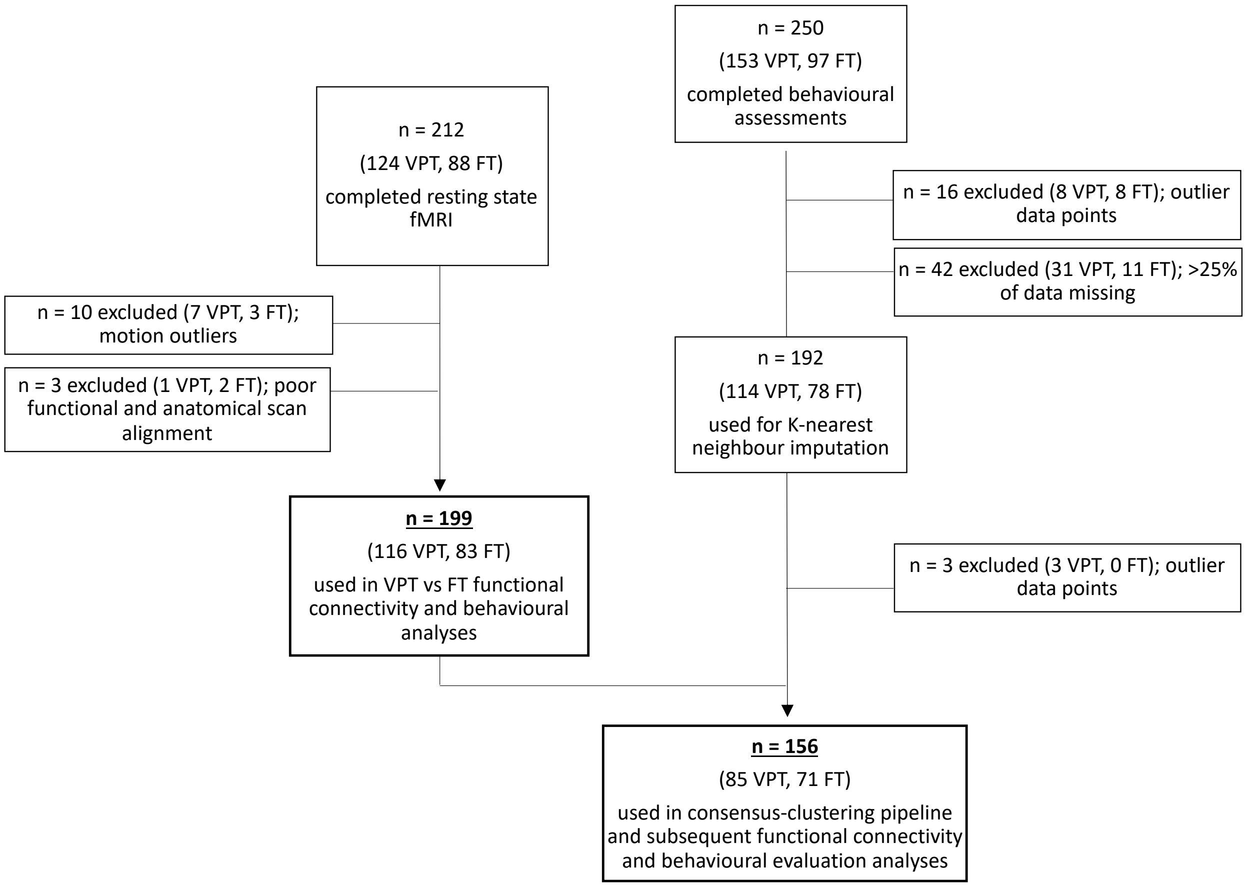


**Figure SM 1. Participants' selection flow diagram.**

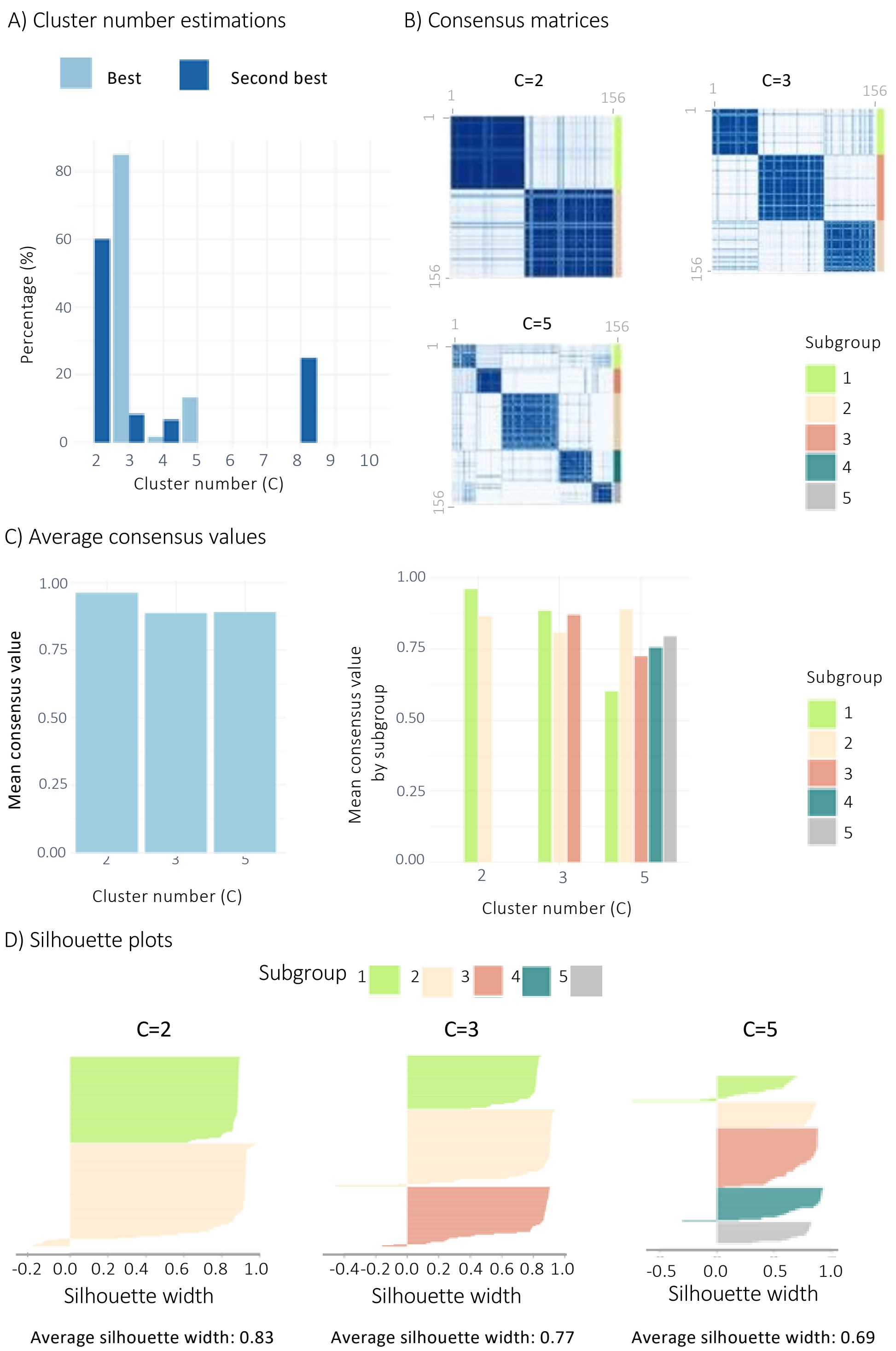


**Figure SM 2. Estimating the optimal number of clusters.** A) Number of times (in percent) each number of clusters was selected as the best (pale blue) or second best (blue) using Eigengap and Rotation Cost for the 30 combinations of K-alpha parameters, with C=2 and C=3 being the most frequently estimated best and second best number of clusters, followed by C=5. B) Consensus matrices from C=2, C=3 and C=5 showing consensus values (i.e., proportion of times each pair of subjects co-clustered into the same cluster over the 1,000 iterations; darker blue indicates higher proportions of co-clustering. C) Mean consensus values for each number of clusters (C=2, C=3 and C=5) and for each subgroup within the different number of clusters runs (left and right, respectively) with the highest values belonging to C=2. D) Silhouette width values for each subgroup within the different number of clusters runs: C=2, C=3 and C=5, whereby C=2 was also displaying the highest values.

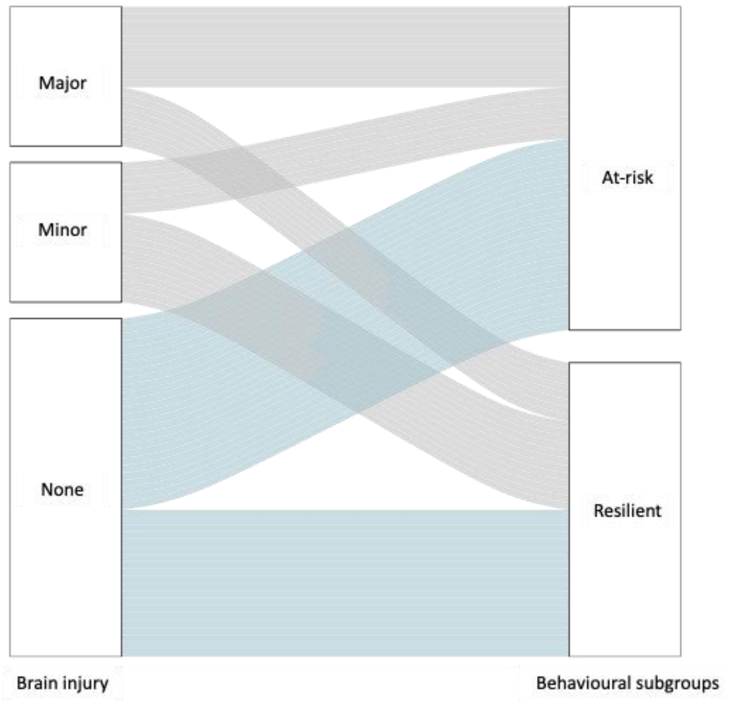


**Figure SM 3. Alluvial plots showing VPT individuals with no brain injury (in blue) and minor or major brain injury (in grey) clustering into the At-risk and Resilient data-driven behavioural subgroups.**

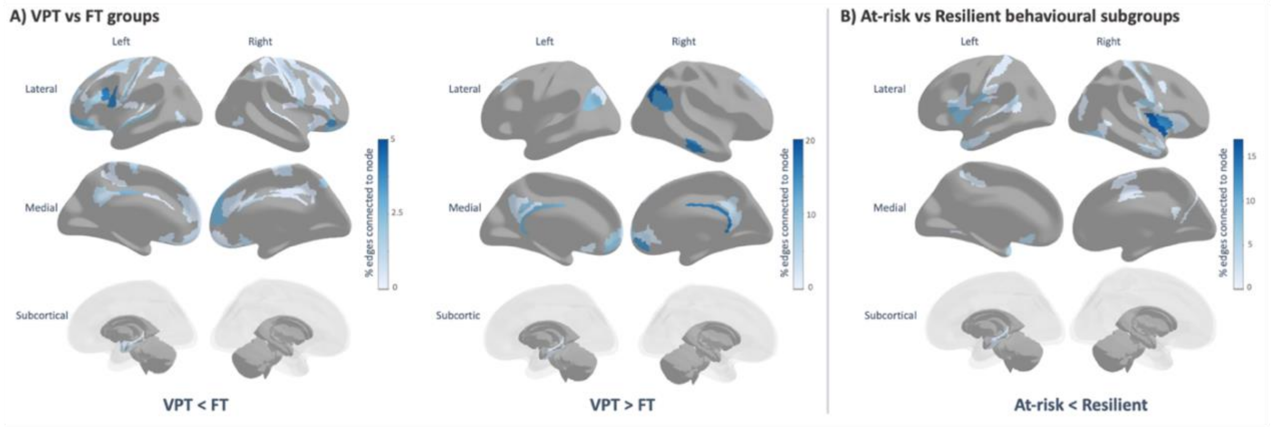


**Figure SM 4. Percentage of edges connected to each region within the significant NBS components at 0.001 p-NBS-Threshold.** Darker colours (blue) denote higher percentages and lighter colours (light blue and white) denote lower percentages, with areas marked in grey indicating regions that are not forming part of the NBS component.

### Supplementary tables

### Table SM 1. Cognitive assessment descriptions.

| Assessment | Assessment description | Construct measured by assessment |
|---|---|---|
| **Hayling Sentence Completion Test (HSCT)** | Participants are asked to complete sentences correctly and incorrectly by providing semantically related or unrelated words. | This task **measures both response initiation speed and response inhibition**. Total scaled score is measured using response latency and response errors. |
| **Controlled Oral Word Association Test (COWAT-FAS)** | Participants are given 60 seconds to list as many words (excluding proper names, numbers, or words with different tenses/endings) as possible beginning with a given letter (i.e., F, A, and S). | This task **measures phonemic verbal fluency**. The total score of phonemic fluency is calculated as the sum of total words produced for each letter. |
| **Stockings of Cambridge (SOC) task** | Participants are required to rearrange three coloured stimuli to match the pattern of the stimuli displayed on the screen in the minimum number of moves possible. | This task **measures spatial planning abilities**. A score of 'Problems Solved in Minimum Moves' is calculated. |
| **Intra-Extra Dimensional Set Shift (IED) task** | Participants are presented with stimuli and are required to learn a rule and select correct responses. The task involves categorising visual stimuli into sets (i.e., the visual discrimination of shapes vs lines) and being able to flexibly respond to changes in stimuli (i.e., shifting attention). The rule changes are intra-dimensional at first (i.e., the shapes or lines are still the relevant stimuli) and then become extra-dimensional (i.e., the shapes or lines are no longer the relevant stimuli). | This task **measures attentional set shifting**. The number of errors made are calculated and adjusted for task stages completed by adding errors for stages not completed. Outcome measure is a 'Total Errors Adjusted' score. |
| **Paired Associates Learning (PAL) task** | Multiple patterns appear on a screen in a random order and then disappear. These patterns reappear, and the participant is asked to recall where on the screen that pattern was originally displayed. | **Measures episodic visuo-spatial associative memory**. The total number of adjusted errors is used a summary score. |
| **Motor Screening Task (MOT)** | A cross appears on the screen and participants are asked to press on it as quickly and as accurately as possible. | This task **measures visuo-motor coordination and comprehension abilities**. The mean reaction time is used as a summary score. |
| **The Trail Making Task – B (TMT-B)** | The task requires participants to draw lines between circles randomly distributed on a paper (whereby each circle contains a letter or a number). They participant is asked to connect the circles sequentially by alternating between numbers and letters. | This task **measures visual scanning, attentional set shifting and cognitive flexibility**. The time required to complete the task is measured and used as a summary score. |

| | | |
|---|---|---|
| **Continuous Performance Test (CPT)** | Participants are presented with a series of continuously changing visual stimuli and the participant is required to respond by pressing a button when a stimulus is presented on the screen and to refrain from responding when a “non-target” stimulus (e.g., “X”) is presented. | This task **measures sustained attention and response inhibition**. The total reaction time for correct responses is used a summary score. |
| **Wechsler Abbreviated Scale of Intelligence (WASI)** | Participants complete a series of tasks measuring visuo-spatial and problem-solving, abstract verbal and non-verbal reasoning, verbal expression, semantic knowledge, and verbal comprehension. | The scaled total IQ score is used as a summary **measure of general intelligence.** |

**Table SM 2. Clinical, socio-demographic, and behavioural profiles of included VPT sample (n=116), relative to VPT excluded from whole sample (n=37).**

| Variable | VPT excluded (n=37) | VPT included (n=116) | p-value |
|---|---|---|---|
| Age at assessment, years | 31.62 (3.39) | 31.37 (3.71) | 0.115 |
| Gestational age at birth, weeks | 28.00 (4.00) | 30.00 (3.00) | 0.024 |
| Birth weight, grams | 1183.00 (253.00) | 1345.00 (512.00) | 0.453 |
| [a] Perinatal brain injury (None: Minor: Major), n | 8:7:13 | 49:22:22 | 0.038 |
| [b] Participants' current socio-economic status (I – II: III: I-V: Student: Unemployed), n | 16: 7: 5: 1: 8 | 51: 41: 6: 1: 16 | 0.094 |
| [b] Parental socio-economic status at birth (I – II: III: I-V), n | 15:9:3 | 43:36:8 | 0.761 |
| Sex M:F, n | 24:13 | 66:50 | 0.506 |
| [c] COWAT, total words | 10.00 (7.25) | 13.00 (5.75) | 0.042 |
| [c] CANTAB – SOC, problems solved | 8.60 (3.00) | 9.00 (2.75) | 0.228 |
| [c] CANTAB – IED, total errors adjusted | 20.60 (41.75) | 15.00 (25.50) | 0.342 |
| [c] TMT-B, time to finish task | 100.50 (51.80) | 73.50 (40.50) | 0.007 |
| [c] CPT, total reaction time for correct responses | 430.90 (84.75) | 417.50 (59.15) | 0.226 |
| [c] WASI – full scale IQ | 100.70 (24.65) | 106 .00(13.75) | 0.097 |
| [c] CANTAB – MOT, reaction time | 717.50 (242.65) | 691.00 (200.80) | 0.990 |
| [d] PDI, total score | 24.50 (43.50) | 21.50 (50.25) | 0.680 |
| [e] AQ10, total score | 3.31 (3.00) | 2.00 (2.44) | 0.031 |
| [f] CAARMS, general psychopathology score | 2.50 (9.25) | 2.00 (5.50) | 0.133 |
| [g] GHQ, total score | 12.00 (6.00) | 10.00 (6.00) | 0.594 |
| [c] ERT, total correct | 53.50 (10.25) | 56.60 (11.15) | 0.092 |

| | | | |
|---|---|---|---|
| **[c] SAS, total score** | 1.49 (0.56) | 1.58 (0.45) | 0.867 |

***Note.*** *Median (interquartile range) reported unless number of participants (n) is reported. [a] Perinatal brain injury rated from ultrasound scans: no haemorrhage (none), grade I – II periventricular haemorrhage without ventricular dilation (minor injury) and grade III – IV periventricular haemorrhage with ventricular dilation (major injury). [b] Socio-economic status occupation classifications. I: Higher managerial, administrative and professional occupations; II: Intermediate occupations, small employers and own account workers; III: Routine and manual occupations – lower supervisory and technical and semi-routine and routine occupations. Missing data for VPT excluded and included respectively: [c](n=9, n=22), [d](n=1, n=22), [e](n=10, n=19), [f](n=1, n=17), [g](n=0, n=9).*

### Table SM 3. VPT only – At-risk and Resilient behavioural subgroup clinical and socio-demographic profiles.

| Variable | VPT Subgroup 1 – Resilient | VPT Subgroup 2 – At-risk | p-value | FDR p-value |
|---|---|---|---|---|
| **Age at assessment, years** | 31.03 (3.34) | 30.50 (3.60) | 0.635 | 0.635 |
| **Gestational age at birth, weeks** | 30.00 (4.00) | 30.00 (3.25) | 0.187 | 0.262 |
| **Birth weight, grams** | 1382.50 (479.50) | 1201.50 (498.75) | 0.035 | 0.081 |
| [a] **Perinatal brain injury, n (%)** | | | 0.303 | 0.416 |
| *None* | 20 (48.78%) | 26 (59.09%) | | |
| *Minor* | 12 (29.27%) | 7 (15.91%) | | |
| *Major* | 8 (19.51%) | 11 (25%) | | |
| [b] **Participants' current socio-economic status, n (%)** | | | | |
| *I – II* | 28 (68.29%) | 16 (36.36%%) | 0.007 | 0.047 |
| *III* | 12 (29.27%) | 17 (38.64%) | | |
| *IV – V* | 0 (0.00%) | 2 (4.55%) | | |
| *Student* | 0 (0.00%) | 1 (2.27%) | | |
| *Unemployed* | 1 (2.44%%) | 8 (18.18%) | | |
| [b] **Parental socio-economic status at birth, n (%)** | | | 0.177 | 0.262 |
| *I – II* | 23 (56.10%) | 16 (36.36%) | | |
| *III* | 12 (29.27%) | 20 (45.46%) | | |
| *IV – V* | 4 (9.76%) | 4 (9.09%) | | |
| **Sex, n (%)** | | | 0.029 | 0.081 |
| *Male* | 30 (73.17%) | 21 (47.73%) | | |
| *Female* | 11 (26.83%) | 23 (52.27%) | | |
| **Total, n** | 41 | 44 | | |

***Note.*** *Median (interquartile range) reported unless stated otherwise where number of participants (n) is reported alongside percentage (%).* [a] *Brain ultrasound scans were used to rate perinatal brain injury into three categories: no haemorrhage (no injury), grade I – II periventricular haemorrhage without ventricular dilation (minor injury) and grade III – IV periventricular haemorrhage with ventricular dilation (major injury).* [b] *Socio-economic status was categorised according to the Office of National Statistics, 1980 occupation classifications. I: Higher managerial, administrative and professional occupations; II: Intermediate occupations, small employers and own account workers; III: Routine and manual occupations – lower supervisory and technical and semi-routine and routine occupations.*

### Table SM 4. FT only – At-risk and Resilient behavioural subgroup socio-demographic profiles.

| Variable | FT Subgroup 1 – Resilient | FT Subgroup 2 – At-risk | p-value | FDR p-value |
|---|---|---|---|---|
| **Age at assessment, years** | 28.82 (3.36) | 29.48 (5.31) | 0.61 | 0.811 |
| **[a] Participants' current socio-economic status, n (%)** | | | 0.035 | 0.138 |
| *I – II* | 18 (60.00%) | 14 (34.15%) | | |
| *III* | 9 (30.00%) | 14 (34.15%) | | |
| *IV – V* | 0 (0.00%) | 0 (0.00%) | | |
| *Student* | 1 (3.33%) | 10 (24.39%) | | |
| *Unemployed* | 1 (3.33%) | 3 (7.31%) | | |
| **[a] Parental socio-economic status at birth, n (%)** | | | 0.217 | 0.435 |
| *I – II* | 21 (70%) | 17 (41.46%) | | |
| *III* | 4 (13.33%) | 10 (24.39%) | | |
| *IV – V* | 1 (3.33%) | 2 (4.88%) | | |
| **Sex, n (%)** | | | 0.831 | 0.831 |
| *Male* | 13 (43.33%) | 20 (48.78%) | | |
| *Female* | 17 (56.67%) | 21 (51.22%) | | |
| **Total, n** | 30 | 41 | | |

***Note.*** *Median (interquartile range) reported unless stated otherwise where number of participants (n) is reported alongside percentage (%).* [a] *Socio-economic status was categorised according to the Office of National Statistics, 1980 occupation classifications. I: Higher managerial, administrative and professional occupations; II: Intermediate occupations, small employers and own account workers; III: Routine and manual occupations – lower supervisory and technical and semi-routine and routine occupations.*

**Table SM 5. Experimenting with p-value thresholds for NBS using two-tailed statistical testing.**

| Two-tailed effect of interest | p-value threshold | Number of identified components | Number of significant components | Significant component size | Significant component strength | FWE p-value |
|---|---|---|---|---|---|---|
| **VPT vs FT** | 0.05 | 1 | 1 | 5244 | 2347.93 | 0.004 |
| | 0.01 | 1 | 1 | 1332 | 509.11 | 0.001 |
| | 0.001 | 22 | 1 | 153 | 50.08 | <0.001 |
| **At-risk vs Resilient** | 0.05 | 1 | 0 | n/a | n/a | n/a |
| | 0.01 | 3 | 1 | 693 | 232.11 | 0.013 |
| | 0.001 | 29 | 1 | 27 | 12.27 | 0.013 |

**Table SM 6. VPT < FT – nodes with the highest number of connections within the significant NBS component.**

| Brain region | HCP-MMP atlas region | Number of edges | Percentage of edges |
|---|---|---|---|
| **Superior temporal gyrus (auditory association cortex)** | right Area STGa | 36 | 2.453988 |
| **Inferior parietal cortex** | left Area PGi | 33 | 2.249489 |
| **Inferior frontal cortex** | right Area 47l (47 lateral) | 32 | 2.181322 |
| **Superior parietal cortex** | left Medial IntraParietal Area | 29 | 1.976823 |
| **Orbitofrontal cortex** | right Area 47m | 27 | 1.840491 |
| **Inferior frontal cortex** | left Area IFJa | 26 | 1.772324 |
| **Orbitofrontal cortex** | left Area anterior 10p | 26 | 1.772324 |
| **Anterior cingulate and medial prefrontal cortex** | left Area p32 | 25 | 1.704158 |
| **Inferior premotor** | left Rostral Area 6 | 25 | 1.704158 |
| **Superior temporal gyrus (auditory association cortex)** | left Area STGa | 25 | 1.704158 |
| **Lateral occipital/posterior temporal visual area** | left Area PH | 25 | 1.704158 |
| **Superior parietal cortex** | left Area Lateral IntraParietal ventral | 24 | 1.635992 |
| **Orbitofrontal cortex** | left Area 47l (47 lateral) | 24 | 1.635992 |
| **Orbitofrontal cortex** | left Area 11l | 24 | 1.635992 |
| **Orbitofrontal cortex** | left Orbital Frontal Complex | 24 | 1.635992 |
| **Lateral temporal cortex** | left Area TE1 anterior | 23 | 1.567825 |
| **Dorsolateral prefrontal cortex** | left Superior Frontal Language Area | 22 | 1.499659 |
| **Inferior frontal cortex** | left Area IFSp | 22 | 1.499659 |

| | | | |
|---|---|---|---|
| **medial temporal lobe** | left ParaHippocampal Area 2 | 22 | 1.499659 |
| **Medial temporal lobe** | right Entorhinal Cortex | 22 | 1.499659 |
| **Lateral temporal cortex** | right Area TG Ventral | 21 | 1.431493 |
| **Posterior cingulate cortex** | left Area 23d | 20 | 1.363327 |
| **Dorsolateral prefrontal cortex** | left Area 8B Lateral | 19 | 1.29516 |
| **Inferior parietal cortex** | left Area PGs | 19 | 1.29516 |
| **Dorsolateral prefrontal cortex** | right Area 8B Lateral | 19 | 1.29516 |
| **Orbitofrontal cortex** | right posterior OFC Complex | 19 | 1.29516 |
| **Dorsal stream visual cortex** | left Ventral Area 6 | 18 | 1.226994 |
| **Anterior cingulate and medial prefrontal cortex** | right Area 9 Middle | 18 | 1.226994 |
| **Inferior parietal cortex** | right Area IntraParietal 2 | 18 | 1.226994 |
| **Orbitofrontal cortex** | right Orbital Frontal Complex | 17 | 1.158828 |
| **Lateral occipital/posterior temporal visual area** | right Area PH | 17 | 1.158828 |
| **Anterior cingulate and medial prefrontal cortex** | right Area 25 | 17 | 1.158828 |
| **Anterior cingulate and medial prefrontal cortex** | left Area 10r | 16 | 1.090661 |
| **Orbitofrontal cortex** | left Area anterior 47r | 16 | 1.090661 |
| **Orbitofrontal cortex** | left Area 13l | 16 | 1.090661 |
| **Orbitofrontal cortex** | right Area 11l | 16 | 1.090661 |
| **Superior temporal gyrus (auditory association cortex)** | right Area TA2 | 16 | 1.090661 |

### Table SM 7. VPT > FT – nodes with the highest number of connections within the significant NBS component.

| Brain region | HCP-MMP atlas region | Number of edges | Percentage of edges |
|---|---|---|---|
| **Posterior opercular cortex** | left Frontal Opercular Area 4 | 30 | 3.118503 |
| **Posterior opercular cortex** | right Frontal Opercular Area 4 | 30 | 3.118503 |
| **Posterior cingulate cortex** | right Complex | 26 | 2.702703 |
| **Inferior parietal cortex** | left Area PF Complex | 24 | 2.494802 |
| **Inferior parietal cortex** | right Area IntraParietal 2 | 24 | 2.494802 |
| **Orbitofrontal cortex** | right Area 10d | 23 | 2.390852 |
| **Inferior parietal cortex** | right Area PGs | 22 | 2.286902 |
| **Posterior opercular cortex** | right Area Frontal Opercular 5 | 22 | 2.286902 |
| **Inferior parietal cortex** | right Area PGi | 20 | 2.079002 |
| **Anterior Cingulate and Medial Prefrontal Cortex** | left Area 10r | 18 | 1.871102 |
| **Posterior cingulate cortex** | right Parieto-Occipital Sulcus Area 1 | 18 | 1.871102 |
| **Dorsolateral prefrontal cortex** | right Area posterior 9-46v | 18 | 1.871102 |
| **Superior temporal gyrus (auditory association cortex)** | right Area STGa | 18 | 1.871102 |
| **Lateral temporal cortex** | right Area TE1 Middle | 18 | 1.871102 |
| **Superior temporal gyrus (auditory association cortex)** | left Area STGa | 16 | 1.663202 |
| **Inferior parietal cortex** | right Area PFm Complex | 16 | 1.663202 |
| **Posterior opercular cortex** | left Area Frontal Opercular 5 | 15 | 1.559252 |
| **Posterior cingulate cortex** | right Area 31pd | 15 | 1.559252 |

| | | | |
|---|---|---|---|
| **Anterior Cingulate and Medial Prefrontal Cortex** | right Anterior 24 prime | 14 | 1.455301 |
| **Anterior Cingulate and Medial Prefrontal Cortex** | right Area a24 | 14 | 1.455301 |
| **Posterior cingulate cortex** | left Area dorsal 23 a+b | 13 | 1.351351 |
| **Posterior cingulate cortex** | left Area 31p ventral | 13 | 1.351351 |
| **Superior parietal cortex (medial)** | left Medial Area 7A | 13 | 1.351351 |
| **Anterior Cingulate and Medial Prefrontal Cortex** | left Area Posterior 24 prime | 13 | 1.351351 |
| **Anterior Cingulate and Medial Prefrontal Cortex** | right Area 10r | 13 | 1.351351 |
| **Dorsolatetal prefrontal cortex** | right Area 8C | 13 | 1.351351 |
| **Posterior opercular cortex** | right Area 43 | 13 | 1.351351 |
| **Posterior cingulate cortex** | left Retrosplenial Complex | 12 | 1.247401 |
| **Inferior parietal cortex** | left Area PGi | 12 | 1.247401 |
| **Superior parietal cortex (medial)** | right Medial Area 7A | 12 | 1.247401 |
| **Anterior Cingulate and Medial Prefrontal Cortex** | right Area 9 Middle | 12 | 1.247401 |
| **Temporo-parietal-occipital junction** | right TemporoParietoOccipital Junction 2 | 12 | 1.247401 |
| **Orbitofrontal cortex** | right Area posterior 10p | 12 | 1.247401 |

**Table SM 8. At-risk < Resilient – nodes with the highest number of connections within the significant NBS component.**

| Brain region | HCP-MMP atlas region | Number of edges | Percentage of edges |
|---|---|---|---|
| **Insular cortex** | right Posterior Insular Area 2 | 32 | 3.846154 |
| **Frontal opercular cortex** | left Area OP4/PV | 25 | 3.004808 |
| **Frontal opercular cortex** | left Frontal Opercular Area 4 | 24 | 2.884615 |
| **Insular cortex** | right Middle Insular Area | 24 | 2.884615 |
| **Frontal opercular cortex** | left Frontal Opercular Area 2 | 23 | 2.764423 |
| **Posterior opercular cortex** | right Area OP4/PV | 23 | 2.764423 |
| **Inferior frontal cortex** | left Area 44 | 22 | 2.644231 |
| **Lateral occipital/posterior temporal visual area** | right Area PH | 21 | 2.524038 |
| **Insular cortex** | left Posterior Insular Area 2 | 20 | 2.403846 |
| **Lateral temporal cortex** | left Area TE2 anterior | 19 | 2.283654 |
| **Superior premotor cortex** | right Dorsal area 6 | 18 | 2.163462 |
| **Lateral temporal cortex** | left Area TG dorsal | 15 | 1.802885 |
| **Posterior opercular cortex** | right Area 43 | 15 | 1.802885 |
| **Temporo-parieto-occipital junction** | left Superior Temporal Visual Area | 14 | 1.682692 |
| **Supplementary motor area** | left Supplementary and Cingulate Eye Field | 14 | 1.682692 |
| **Insular cortex** | left Insular Granular Complex | 14 | 1.682692 |
| **Lateral temporal cortex** | right Area TF | 14 | 1.682692 |
| **Posterior opercular cortex** | left Area OP2-3/VS | 13 | 1.5625 |
| **Anterior Cingulate and Medial Prefrontal Cortex** | left Area 25 | 13 | 1.5625 |

| | | | |
|---|---|---|---|
| **Primary somatosensory cortex** | right Primary Sensory Cortex | 13 | 1.5625 |
| **Posterior opercular cortex** | right Area OP1/SII | 12 | 1.442308 |
| **Insular cortex** | right Area 52 | 12 | 1.442308 |
| **Primary somatosensory cortex** | left Primary Sensory Cortex | 11 | 1.322115 |
| **Insular cortex** | left Middle Insular Area | 11 | 1.322115 |
| **Frontal opercular cortex** | left Frontal Opercular Area 3 | 11 | 1.322115 |
| **Superior temporal sulcus (auditory association cortex)** | left Area STSd anterior | 11 | 1.322115 |
| **Lateral temporal cortex** | left Area TF | 11 | 1.322115 |
| **Anterior Cingulate and Medial Prefrontal Cortex** | right Area 8BM | 11 | 1.322115 |
| **Posterior opercular cortex** | right Area OP2-3/VS | 11 | 1.322115 |
| **Frontal opercular cortex** | right Frontal Opercular Area 3 | 11 | 1.322115 |
| **Posterior opercular cortex** | left Area PFcm | 10 | 1.201923 |
| **Posterior opercular cortex** | left Frontal Opercular Area 1 | 10 | 1.201923 |
| **Frontal opercular cortex** | right Frontal Opercular Area 2 | 10 | 1.201923 |
| **Superior temporal sulcus (auditory association cortex)** | right Area STSd anterior | 10 | 1.201923 |
| **Lateral temporal cortex** | right Area TE1 anterior | 10 | 1.201923 |

**Table SM 9. Sensitivity analyses – NBS component results at p-NBS-threshold = 0.001.**

| | **Edges, n (% of all possible connections)** | **Nodes, n (% of all regions)** | **Component strength, T-stat** | **FWE p-value** |
|---|---|---|---|---|
| ***VPT < FT*** | 221 (0.32%) | 179 (47.86%) | 77.89 | 0.001 |
| ***VPT > FT*** | 42 (0.060%) | 29 (7.75%) | 16.36 | 0.001 |
| ***At-risk < Resilient*** | 60 (0.086%) | 52 (13.90%) | 24.84 | 0.006 |
